# Molecular mechanism of parental H3/H4 recycling at a replication fork

**DOI:** 10.1101/2024.03.07.583824

**Authors:** Fritz Nagae, Yasuto Murayama, Tsuyoshi Terakawa

## Abstract

In eukaryotic chromatin replication, faithful recycling of histones from parental DNA to replicated leading and lagging strands is essential for maintaining epigenetic information across generations. A recent experimental study has revealed that disrupting interactions between the N-terminal disordered tail of Mcm2, a subunit in DNA replication machinery, and a histone H3/H4 tetramer, a carrier of epigenetic modifications, perturbs their faithful recycling. However, the molecular pathways via which the H3/H4 tetramer attached to Mcm2 is recycled to the replicated strands have yet to be deciphered. Furthermore, the factors that regulate the ratio recycled to each strand and the destination location still need to be discovered. The elucidation requires visualization of a structural trajectory from H3/H4 bound to Mcm2 until recycled to the replicated strands. In this study, we performed molecular dynamics simulations of yeast DNA replication machinery (Mcm2-7, Cdc45, GINS, Pol ε, and RPA), an H3/H4 tetramer, and replicated DNA strands. The simulations demonstrated that histones are recycled via Cdc45-mediated and unmediated pathways even without histone chaperones, as our *in vitro* biochemical assays supported. Also, RPA binding to the ssDNA portion of the lagging strand regulated the ratio recycled to each strand, whereas DNA bending by Pol ε modulated the destination location. Together, the simulations in this study provided testable hypotheses, which are vital for clarifying intracellular histone recycling controlled by the cooperation of many histone chaperones.

## INTRODUCTION

Eukaryotic DNA forms chromatin, a linear array of nucleosomes, each composed of one H3/H4 tetramer and two H2A/H2B dimers wrapped by 147 base-pairs (bp) DNA^1^. These nucleosomes modulate DNA transactions such as DNA replication, transcription, and repair by permitting or occluding access of regulatory proteins to DNA depending on histone variants or histone post-translational modifications^2,3^. Therefore, nucleosomes and their constituent histones carry epigenetic information that regulates DNA transactions^3–5^. In chromatin replication, DNA replication machinery (replisome) composed of Mcm2-7, Cdc45, GINS, Pol ε, and RPA inevitably collides with a nucleosome, which must be dismantled to allow the helicase to pass through the nucleosome array^6–8^. Subsequently, the dismantled histones are recycled to replicated leading or lagging strands^6–10^. Faithful histone recycling is vital for maintaining epigenetic information across generations^4,5,11–13^.

A recent experimental study revealed that disrupting the interaction between the N-terminal intrinsically disordered tail (N-tail) of Mcm2, a subunit of Mcm2-7, and a histone H3/H4 tetramer perturbs faithful histone recycling^14^. Indeed, the Mcm2 N-tail interacted with the H3/H4 tetramer in the crystal structure^15,16^. Also, previous *in vitro* biochemical assays showed that the Mcm2 N-tail promotes nucleosome assembly on DNA^17^, suggesting that the Mcm2 N-tail directly deposits the H3/H4 tetramer to the replicated strands. As a support, these interactions are essential for preserving heterochromatin silencing at sub-telomeric loci^18^. Recent experimental studies also demonstrated that nucleosomes were assembled on the replicated strands in the presence of histone chaperones and chromatin remodelers^19,20^. However, it remains unknown whether the Mcm2 N-tail alone is sufficient to hand over the H3/H4 tetramer to the replicated strands or whether additional histone chaperones are necessary. Also, the molecular pathways via which the H3/H4 tetramer attached to Mcm2 is recycled to the replicated strands have yet to be deciphered.

Symmetric histone recycling to the two replicated strands underlies the maintenance of cellular identity after cell division^4,14,18,21–23^. In contrast, asymmetric histone recycling alters gene expression profiles in the two daughter cells and potentially triggers cell differentiation^4,5,11,24^. Therefore, the mechanism to regulate the ratio recycled to each strand is vital for organisms to maintain or change their cellular state. Also, the destination location may fine-tune the gene expression or necessitate dramatic nucleosome remodeling after chromatin replication. However, the factors that regulate the ratio recycled to each strand and the destination location still need to be discovered.

Hitherto, deep sequencing of chromatin in nucleus revealed the ratio recycled to each strand and the nucleosome position after chromatin replication *in vivo*^14,21,22,25,26^. However, since various biomolecules are mixed in the nucleus, it has been difficult to clarify the molecular mechanism by which minimal factors achieve histone recycling. An *in vitro* experiment was also performed to reconstitute the recycling reaction with only purified components^19^. However, the study did not analyze the ratio and the destination location. Previous cryo-electron microscopy studies determined the structures of the DNA replication machinery engaging the replicated DNA^27–30^. However, these static structures lack flexible regions, including the Mcm2 N-tail and the replicated leading and lagging DNA strands. Also, it has been challenging to elucidate the dynamics of the H3/H4 recycling. Overcoming these challenges requires visualization of the molecular structural trajectory from H3/H4 bound to Mcm2 until handed over to the replicated strands.

In this study, we first performed coarse-grained molecular dynamics simulations of yeast DNA replication machinery containing Mcm2-7, Cdc45, GINS, Pol ε, and RPA, bound to an H3/H4 tetramer, and replicated DNA strands. Previous studies carefully calibrated the coarse-grained model to reproduce the intrinsically disordered tail dynamics^31^, electrostatic interactions^32^, and nucleosome assembly dynamics^33,34^, all required for the current histone recycling simulations. In this study, we also calibrated the interaction parameters of the Mcm2 N-tail and the H3/H4 tetramer. The simulations demonstrated that H3/H4 tetramers can be recycled to replicated strands without histone chaperones, as supported by *in vitro* replication assays using purified proteins in the current study. The simulation trajectories also revealed two dominant pathways for histone recycling: the Cdc45-mediated and unmediated pathways. In the Cdc45-mediated pathway, the H3/H4 tetramer is once bound to Cdc45 and handed over to the leading strand. On the other hand, in the Cdc45-unmediated pathway, the tetramer is directly handed over to the lagging strand without binding to Cdc45. Consistent with the Cdc45-mediated pathway, the native-polyacrylamide gel electrophoresis (Native-PAGE) assays confirmed that Cdc45 and H3/H4 tetramer electrostatically interact with each other. Also, RPA binding to the ssDNA portion of the lagging strand regulated the ratio recycled to each strand, whereas DNA bending by Pol ε modulated the destination location. Together, the simulations provided valuable insights and experimentally verifiable hypotheses concerning the molecular mechanism *in vitro*, which is crucial for elucidating the mechanism of *in vivo* histone recycling regulated by the collaborative actions of multiple histone chaperones.

## RESULTS

### Modeling of the replicated-DNA-engaged replisome binding to the H3/H4 tetramer

Molecular structural models and potential energy functions for protein and DNA are required to perform molecular dynamics simulations of the replicated-DNA-engaged replisome binding to the H3/H4 tetramer. This study adopted the AICG2+ model for proteins (see the original paper^35^ for details) in which one particle at the C_α_ atom represents one amino acid. The original study optimized the parameters of the potential energy functions, which stabilizes the reference native protein structure, to reproduce fluctuations around the structure^35^. Also, the parameters derived from feature statistics of the loop structures registered in the Protein Data Bank (https://www.rcsb.org/) allow the model to reproduce the structural ensemble of the disordered regions^31^. We also adopted the 3SPN.2 model for DNA (see the original paper^36^ for model details). In this model, one nucleotide is represented by three particles placed at the base, sugar, and phosphate sites. The original study optimized the parameters of the potential energy functions, which stabilize the B-form DNA structure, to reproduce the persistence length of both single-(ss) and double-stranded (ds) DNA, the melting temperature, and the hybridization rate. These features, especially the ability to reproduce structural ensembles of disordered regions of proteins and the persistence length of DNA, are essential for the current simulations.

Potential energy functions for protein-protein and protein-DNA interactions model excluded volume and electrostatics. As for the electrostatics, charges were arranged on the protein surface residues based on the RESPAC algorithm^32^ to reproduce the peripheral electrostatic potential of all-atom structures. In addition, the charges of phosphate groups of DNA were set to −0.6 for intra-DNA interactions to model the counter ion condensation around the phosphate groups within the framework of the Debye-Hückel model^36^. On the other hand, the phosphate charges were set to −1.0 for protein-DNA interactions to account for releases of counter ions upon association of proteins to DNA. These potential energy functions have successfully reproduced the dynamics of the bacterial architectural protein HU^37^, the histone chaperone Nap1^38^, and the DNA mismatch recognition protein MutS along DNA^39^ in previous studies. To model the interaction between the Mcm2 N-tail and the H3/H4 tetramer, we added a function (a so-called Gō-like potential) stabilizing the reference native protein structure^15^. The parameters of this potential play a decisive role in H3/H4 competition between Mcm2 N-tail and replicated strands in histone recycling. Therefore, in this study, we performed temperature replica exchange simulations of Mcm2 and an H3/H4 tetramer associating to and dissociating from each other with varying parameters and selected the one that reproduced the experimental binding free energy (Supplementary Figure 1) (Simulation: −10.00 ± 0.26 kcal/mol, Experiment: −10.45 ± 0.04 kcal/mol^15^). To model the interaction between the H3/H4 tetramer and DNA, we added a hydrogen bonding potential as in the previous studies that successfully recapitulated nucleosome stability and DNA unwrapping dynamics^33,34^.

To prepare the initial structure of the CMG helicase, we used the cryo-EM structure (PDB ID: 6U0M^27^) as a reference. The CMG helicase comprises three proteins: Cdc45, Mcm2-7, and GINS^40^ (Figure 1A). Cdc45 is a protein consisting of a single subunit. Mcm2-7 is a protein consisting of six subunits: Mcm2/3/4/5/6/7. GINS is a protein consisting of four subunits: Sld5, Psf1, Psf2, and Psf3. For brevity, we omitted the intrinsically disordered N- and C-termini of the Mcm3/4/5/6/7 subunits. Missing regions in the Cryo-EM structure were treated as intrinsically disordered regions (IDRs) as in the other proteins.

**Figure 1:**
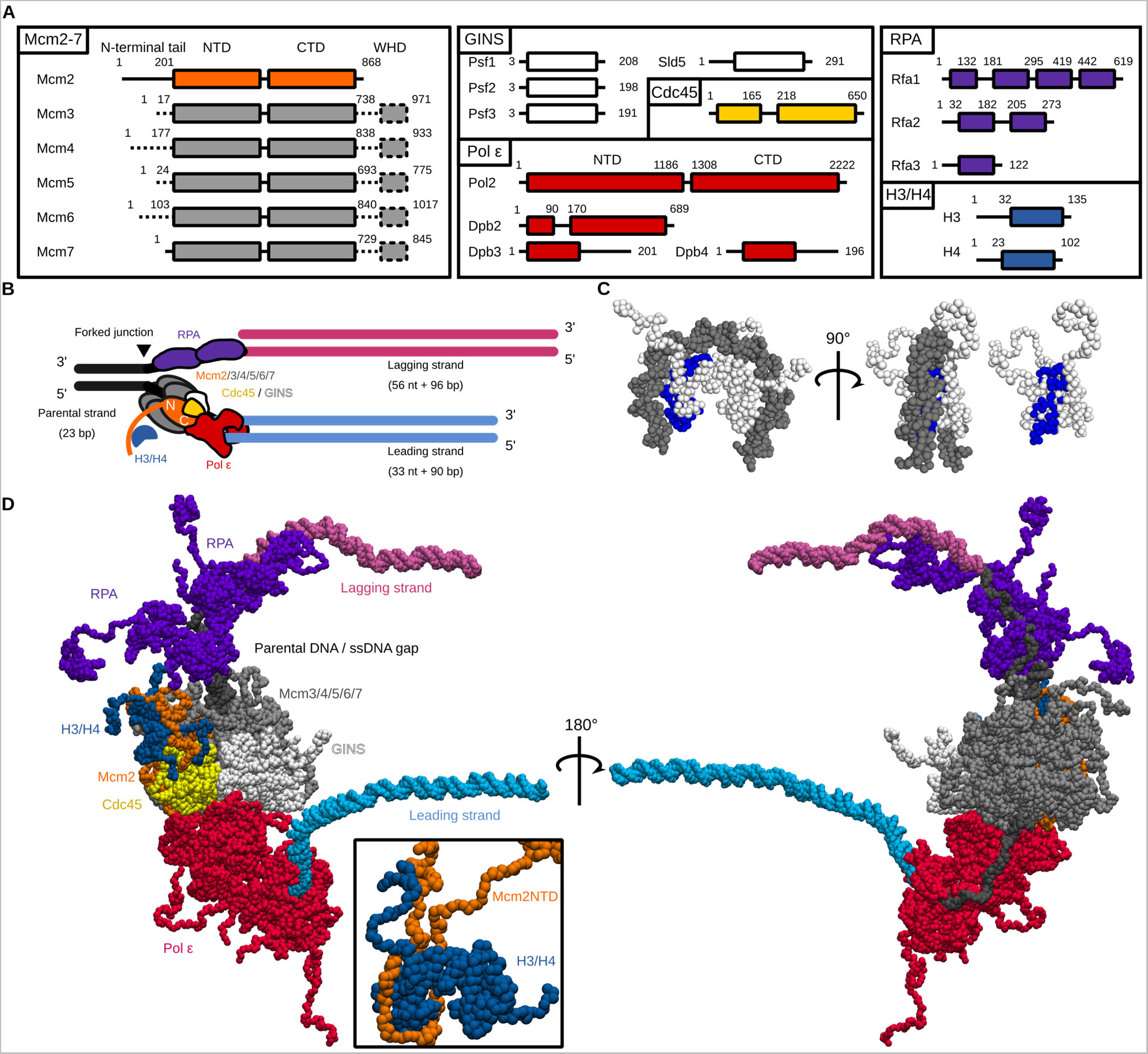
The initial structure for the coarse-grained molecular dynamics simulations of a replicated-DNA-engaged replisome binding to an H3/H4 tetramer. **(A)** The domain composition of Mcm2-7, GINS, Cdc45, Pol ε, RPA, and the H3/H4 tetramer. The dotted lines represent the regions not included in the simulations. **(B)** Schematic illustration of the initial structures. The black, pink, and cyan lines represent the parental, lagging, and leading strands. The orange, gray, white, yellow, red, blue, and purple objects represent Mcm2, Mcm3/4/5/6/7, GINS, Cdc45, Pol ε, H3/H4, and RPA, respectively. **(C)** The structures of a tetrasome and an H3/H4 tetramer. The residues contacting Mcm2 in the crystal structure (PDB ID: 4UUZ) are colored blue. **(D)** The initial coarse-grained structures of the replicated-DNA-engaged replisome.

For Pol ε, we used the cryo-EM structure (PDB ID: 6HV9^28^) as a reference. Pol ε comprises four proteins: Pol2, Dpb2, Dpb3, and Dpb4^41^ (Figure 1A). The reference structure contains part of Pol2 (residues 1308-2222) and Dpb2 (residues 1-90) interacting with the CMG helicase. To obtain the full-length Pol ε reference structure, we superimposed the cryo-EM structure of the entire Pol ε holoenzyme (PDB ID: 6WJV^41^) and Dpb2 (PDB ID: 6HV9^28^) to the reference structure of the CMG helicase (PDB ID: 6U0M^27^).

RPA, which binds to ssDNA gaps to protect it from nucleolytic degradation^42,43^, is a protein consisting of three subunits: Rfa1, Rfa2, and Rfa3 (Figure 1A). The reference structure of Rfa1 comprises the partial structures of DBD-F (PDB ID: 5OMB^44^), DBD-A (PDB ID: 1YNX^45^), DBD-B (PDB ID: 1JMC^46^, homology model), and DBD-C (PDB ID: 6I52^47^). The reference structure of Rfa2 comprises the partial structures of the winged helix domain (PDB ID: 4OU0^48^, homology model) and DBD-D (PDB ID: 6I52^47^). The reference structure of Rfa3 comprises the structure of DBD-E (PDB ID: 6I52^47^). The structures of Rfa1, Rfa2, and Rfa3 were assembled by superimposing DBD-C of Rfa1, DBD-D of Rfa2, and DBD-E of Rfa3 to the heterotrimeric cryo-EM structure (PDB ID: 6I52^47^).

For the H3/H4 tetramer, we used the crystal structure of a nucleosome core particle (PDB ID: 1KX5^49^) as the reference structure. The model was the same as that previously employed to study the nucleosome dynamics upon a collision with a DNA translocase in the presence^38^ and absence^50^ of a histone chaperone.

We modeled the replicated DNA strands based on the DNA conformation in the cryo-EM structure (PDB ID: 6U0M^27^) (Figure 1B). The DNA comprises 11 bp parental dsDNA and 12 nt leading ssDNA. Then, we extended the parental dsDNA region to 23 bp, corresponding to the typical length of linker DNA between nucleosomes in budding yeast genome^51,52^. The missing nucleotides in the leading ssDNA were filled, and the lagging ssDNA region was extended to 56 nt by superimposing the ideal B-form ssDNA structures. The leading and lagging dsDNA region was extended to 90 and 96 bp, sufficient to wrap around the H3/H4 tetramer to form a tetrasome by superimposing the ideal B-form dsDNA structures.

To assemble all the components, we manually placed the H3/H4 tetramers and the two RPA molecules proximal to the Mcm2 N-tail and the 59 nt lagging ssDNA region^53^, respectively, and performed the equilibration molecular dynamics simulation for 1 × 10^6^ steps so that these molecules associate with their binding site. The Mcm2 N-tail bound to the H3/H4 tetramer (residues 1–200) as in the crystal structure (PDB ID: 4UUZ^15^) and occupied the DNA binding interface of the H3/H4 tetramer (Figure 1C). Notably, previous single-molecule fluorescence imaging^54^ showed the two RPA molecules at the replication fork in the physiological concentration of DNA polymerase α (Pol α)^55^. We used the assembled structure as the initial structure for the production molecular dynamics simulations (Figure 1D).

Starting from this initial structure, we performed Langevin dynamics simulations using the software CafeMol 3.2 (https://www.cafemol.org)^56^. The integration timestep was 0.3 CafeMol time units (~14.7 fs). Temperature and friction constant were set to 300K and 0.843, respectively. The monovalent ion concentration of the Debye-Hückel model was set to 300 mM, and the dielectric constant was set to 78.0. These parameters are the same as those of the previous study in which nucleosome stability and DNA unwrapping dynamics in a physiologically relevant condition were successfully recapitulated^33,34^.

Replisome directly recycles an H3/H4 tetramer to replicated DNA strandsWe performed 100 runs of the simulation of a replicated-DNA-engaged replisome binding to an H3/H4 tetramer for 1 × 10^8^ steps, roughly corresponding to ~1.5 microseconds. In 20% (20/100) of the simulation trajectories, the Mcm2 N-tail directly deposited the H3/H4 tetramer onto either of the two replicated strands (Figure 2A & 2B, MovieS1–S4). The other simulation trajectories showed the association of the H3/H4 tetramer with the parental strand (26/100, Supplementary Figure 2A) or no association (54/100). We estimated that it would take ~1 × 10^9^ steps for the H3/H4 tetramer to associate with the parental, leading, or lagging strand in all the trajectories (Supplementary Figure 2B). The H3/H4 tetramer did not dissociate from DNA during the simulations once it was deposited (Figure 2C, Supplementary Figure 2C), indicating that the tetramer deposited on the parental strand (26/100, Supplementary Figure 2B) may require additional factors such as histone chaperones to be evicted. Together, the simulation trajectories suggested that replisome can directly recycle an H3/H4 tetramer to replicated strands in a microsecond timescale, even without histone chaperones.

**Figure 2:**
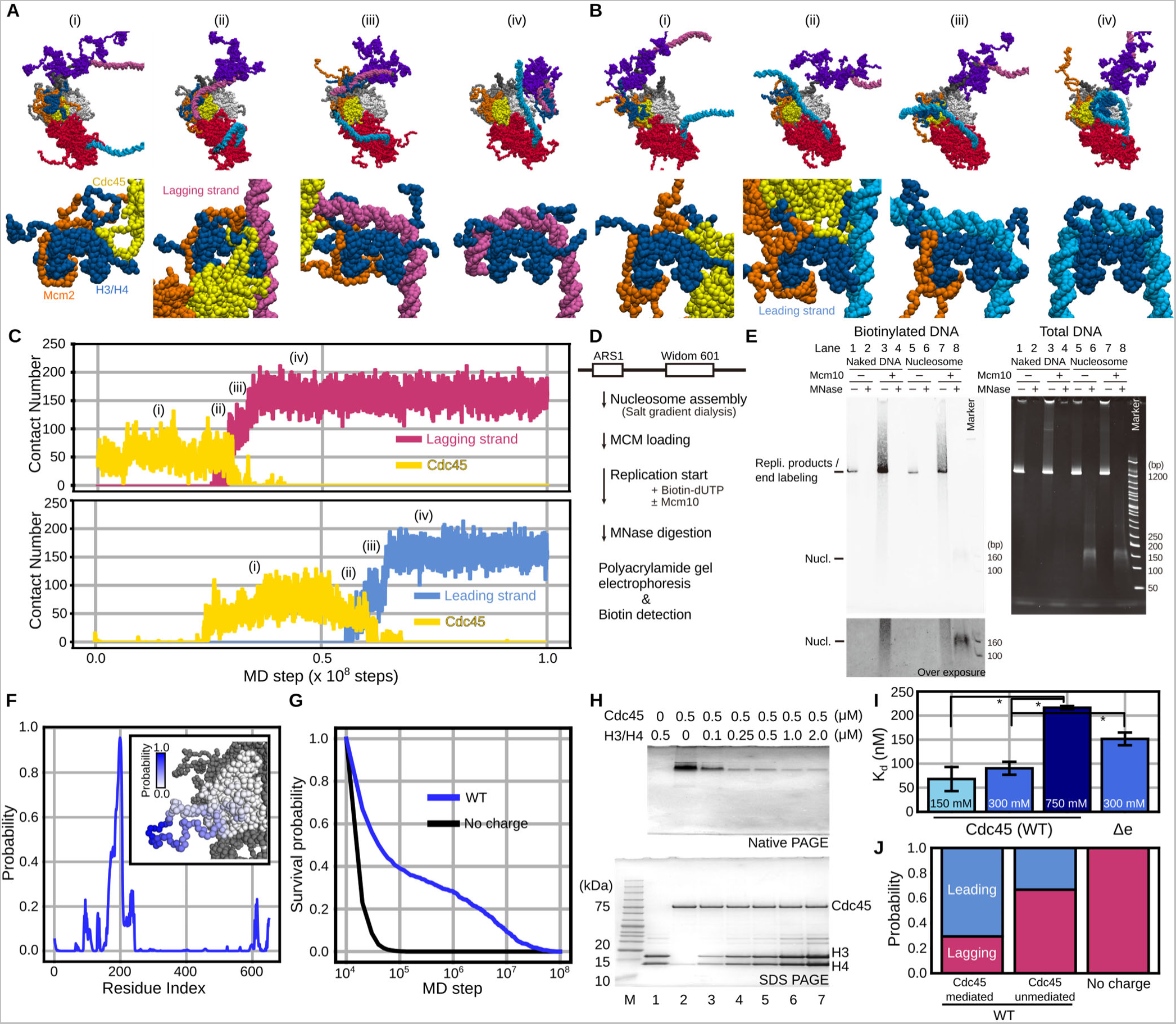
The H3/H4 tetramer was recycled to the replicated strands by the Mcm2 N-tail in the simulations of a replicated-DNA-engaged replisome. **(A, B)** Representative snapshots of the simulation trajectories in which the H3/H4 tetramer was deposited on the lagging (A) and leading (B) strands. The top panels are the magnified version of the bottom ones. **(C)** Time trajectories of the number of residues in the H3/H4 tetramer contacting the lagging strand (pink in the top panel), the leading strand (cyan in the bottom panel), and Cdc45 (yellow) from the trajectories in (A) and (B). **(D)** Schematic of the *in vitro* DNA replication with nucleosome-assembled DNA. **(E)** Gel images of biotinylated DNA (left) and total DNA (right) before and after MNase digestion on native gels. **(F)** Probability of each residue in Cdc45 contacting the H3/H4 tetramer in the simulations. The inset figure is the probabilities represented by shades of blue on the Cdc45 structure. **(G)** Survival probabilities of the association between Cdc45 and the H3/H4 tetramer in the simulations in the presence (WT) or absence (No charge) of charges in the Cdc45 acidic loop. **(H)** Gel images of native PAGE (top) and SDS-PAGE (bottom) of the mixtures of Cdc45 and H3/H4 tetramer (lanes 1-7). ‘M’ denotes a marker (ATTO; 2332346). **(I)** Plots of the intensities of the Cdc45 band in the native gel normalized by the intensity of the band in the absence of H3/H4 tetramer. The normalized intensities of the Cdc45-7HIS (WT) band in different salt concentrations (left). The normalized intensities of the Cdc45-7HIS (WT) and Cdc45-7HIS (D2N, E2Q) bands in 300 mM KCl concentration (right). **(J)** Ratios of the replicated strands to which the H3/H4 tetramer was recycled.

Next, we sought to experimentally confirm whether a replisome can recycle histones to replicated DNA strands upon collision with a nucleosome without histone chaperones. For this purpose, we purified the budding yeast histones and the replication-related proteins to biochemically reconstitute DNA replication^57–59^ with nucleosome-assembled DNA. We prepared the 1155 bp linear DNA substrate containing Autonomously Replicating Sequence 1 (ARS1) and the Widom 601 sequence (Figure 2D, Supplementary Table S1). The nucleosome was initially reconstituted on the DNA substrate by the salt gradient dialysis method^60^, and then the CMG helicases were assembled on the substrate to initiate the replisome-dependent DNA replication. Nascent DNA syntheses were labelled by incorporating biotinylated deoxy uridine nucleotide (biotin-dUTP). After incubation for sufficient time (20 minutes) for replication to complete (Figure 2E left, lanes 3 and 7), we treated the reactions with micrococcal nuclease (MNase). This digests nucleosome-free DNA regions, whereas the nucleosome-coated DNA segments were protected from the digestion, generating ~150 bp DNA fragments. The products were analyzed by polyacrylamide gel electrophoresis. We observed ~150 bp biotinylated DNA fragments in the reaction performed with the nucleosome substrate, whereas no detectable signal was seen when naked DNA was used as a substate (compare lanes 4 and 8 in Figure 2E left). This observation suggested that nucleosomes were formed (recycled) on the replicated DNA strands. However, this ~150 bp band could be generated by end-labeling of the MNase-digested, non-replicated nucleosomal DNA by Pol ε rather than nucleosome reassembly during the replisome-dependent DNA replication. To rule out the possibility, we repeated the assay without Mcm10, which is essential for DNA replication initiation by replisomes. In this case, no detectable biotinylated ~150 bp band was seen (compare lanes 6 and 8 in Figure 2E left). On this gel (Figure 2E right) on which both pre-replicated and post-replicated DNA can be detected, we also observed a similar level of nucleosome assemblies both in the presence and absence of Mcm10, demonstrating that the biotinylated ~150 bp band was not majorly produced by end-labeling. Together, these biochemical assays supported the simulation prediction that a replisome can recycle histones to replicated DNA strands upon collision with a nucleosome without histone chaperones.

### An H3/H4 tetramer is recycled via two pathways

Interestingly, in all the trajectories (100/100), we found that the H3/H4 tetramer bound to the Mcm2 N-tail associated with Cdc45 at least once before being deposited to replicated DNA (Figure 2C, Supplementary Figure 2C & 2D). From the structural point of view, half of the DNA binding surface of the H3/H4 tetramer was wrapped by the Mcm2 N-tail, and the other half was exposed to solvent (Figure 1C & 1D). In the simulation trajectories, this exposed surface was associated with Cdc45 [(i) in Figure 2A–2C]. In the recycling trajectories (20/100), either of the leading or lagging strand fluctuated around the Cdc45-associated H3/H4 tetramer, competed for the binding surface on the H3/H4 tetramer with Cdc45 [(ii) in Figure 2A–2C], and took it away [(iii) in Figure 2A–2C], completing recycling. We calculated the probabilities that each residue of Cdc45 contacts the H3/H4 tetramer and found that residues 184–205 of Cdc45 frequently contacted the H3/H4 tetramer (Figure 2F). These Cdc45 residues are in the flexible acidic loop not resolved in the cryo-EM structures and associated with the H3/H4 tetramer for ~9 × 10^6^ steps on average (Figure 2G). As expected from the hypothesis that the interaction between the Cdc45 acidic loop and H3/H4 tetramer is mainly electrostatic interactions, we could hardly observe contacts lasting longer than 1 × 10^5^ steps between Cdc45 and the H3/H4 tetramer when the simulations were repeated without the charge in the Cdc45 acidic loop (Figure 2G). After the binding, in 85% (17/20) of the recycling trajectories, the leading or lagging strand took the H3/H4 tetramer away from Cdc45 (Movie S1 & S2). In the others (3/20), the Mcm2 N-tail directly deposited the H3/H4 tetramer free from Cdc45 on the leading or lagging strands (Movie S3 & S4, Supplementary Figure 2C–2F). Thus, the simulations showed that the H3/H4 tetramer can be recycled to the replicated strand via the Cdc45-mediated and unmediated pathways.

Next, we sought to perform Native-PAGE assays to experimentally confirm that Cdc45 associates with an H3/H4 tetramer. Thus, we reconstituted H3/H4 tetramers with recombinant histones and purified the budding yeast Cdc45 from *E.coli* as described previously^61^ (Supplementary Figure S3A). We mixed the purified Cdc45 molecules and the reconstituted H3/H4 tetramers, incubated the mixtures for 15 minutes at 30℃, and ran the reaction products on native and denaturing gels. In this assay, the H3/H4 tetramers did not enter the native gels due to their high positive net charge (Figure 2H, lane 1). Interestingly, the intensity of the Cdc45 band (Figure 2H, lane 2) gradually decreased as the concentration of the H3/H4 tetramers increased (Figure 2H, lane 3-7). This is thought to be because Cdc45 no longer enters the gel when it forms a complex with an H3/H4 tetramer due to the high positive net charge of the complex (+40e). Therefore, this result suggested that Cdc45 associates with an H3/H4 tetramer, which is consistent with the simulations.

To examine the contribution of electrostatic interactions to the binding, we repeated the assay with varying KCl concentration (Supplementary Figure S3B). Then, we measured the apparent dissociation constants, which were 68 ± 25 nM, 90 ± 14 nM, and 216 ± 4 nM in 150 mM, 300 mM, and 750 mM KCl concentration, respectively (Figure 2I). The stable association even in 750 mM KCl suggested that hydrophobic interactions, which was not modelled in the simulations, also contributes to the complex formation. Nevertheless, This result suggested that electrostatic interactions significantly contribute the interaction between Cdc45 and an H3/H4 tetramer in concordance with the simulations.

To more specifically confirm that the acidic loop in Cdc45 contributes to the binding to an H3/H4 tetramer, we repeated the assay with a mutant (Δe) in which all the aspartic and glutamic residues were replaced with the asparagine and glutamine residues, respectively (Supplementary Figure S3C). Then, we measured the apparent dissociation constants, which were 90 ± 14 nM and 152 ± 13 nM for the wild-type and mutant proteins, respectively (Figure 2I). This result suggests that electrostatic interactions between the Cdc45 acidic loop and the H3/H4 tetramer contribute to the complex formation in accordance with the simulations.

Next, we sought to analyze the ratio recycled to each strand in the Cdc45-mediated and -unmediated pathways. Thus, we classified the trajectories based on the recycling pathway and the strand to which the H3/H4 tetramer was deposited. In the trajectories with the Cdc45-mediated pathway, 75% (12/16) and 25% (4/16) of them showed deposition to the leading and lagging strands, respectively (Figure 2J, Movie S1 & S2). On the other hand, in the trajectories with the Cdc45-unmediated pathway, 25% (1/4) and 75% (3/4) of them showed deposition to the leading and lagging strands, respectively (Figure 2J, Movie S3 & S4). To confirm the effect of Cdc45 binding to the H3/H4 tetramer on the strand bias, we eliminated electrostatic interactions between Cdc45 and the H3/H4 tetramer by neutralizing the charge of the Cdc45 acidic loop and performed 50 runs of simulations. Of them, only two trajectories (2/50) showed successful histone recycling, both resulting in deposition to the lagging strand (Figure 2J). These results support that the association between Cdc45 and the H3/H4 tetramer promotes histone recycling, especially to the leading strand.

In histone recycling, the leading or lagging strand takes the H3/H4 tetramer away from the Mcm2 N-tail. Thus, the extent to which the leading and lagging strands probe the H3/H4 tetramer bound to the Mcm2 N-tail by their conformational fluctuations is critical for successful recycling. To investigate this range, we defined a vector from the center of mass (COM) of Mcm2-7 to the end of the leading or lagging strand. Then, we calculated the angle between the vector and the longitudinal (elevation angle φ) or lateral (azimuthal angle θ) axis of Mcm2-7. By definition, elevation and azimuthal angles are zero when the vector orients to the direction from the COM of Mcm2-7 to Mcm2 (Figure 3A & 3B). We assumed that the dsDNA bending only negligibly affects the analysis since the strand length (~90 bp) is shorter than the persistence length (~150 bp). The analysis showed that the lagging strand tended to orient to the direction with an elevation angle of 4° ± 36° and an azimuthal angle of 160° ± 79° (Figure 3C). The steric hindrance between the CMG helicase complex and the lagging strand explains the slightly positive mean elevation angle (Figure 3D). Also, the binding of the lagging strand to the CMG helicase explains why the strand was oriented away from Mcm2. Indeed, the lagging strand was associated with the Mcm3, Mcm5, and Mcm7 zinc finger domains in the simulations (Supplementary Figure 4) in line with the cryo-EM structure^27^. On the other hand, the leading strand tended to orient to the direction with an elevation angle of 35° ± 23° and an azimuthal angle of 67° ± 52° (Figure 3E). The association between the leading strand and the N-terminal domain of Pol2, which bent DNA and fixed its orientation, explains this tendency (Figure 1D & 3F).

**Figure 3:**
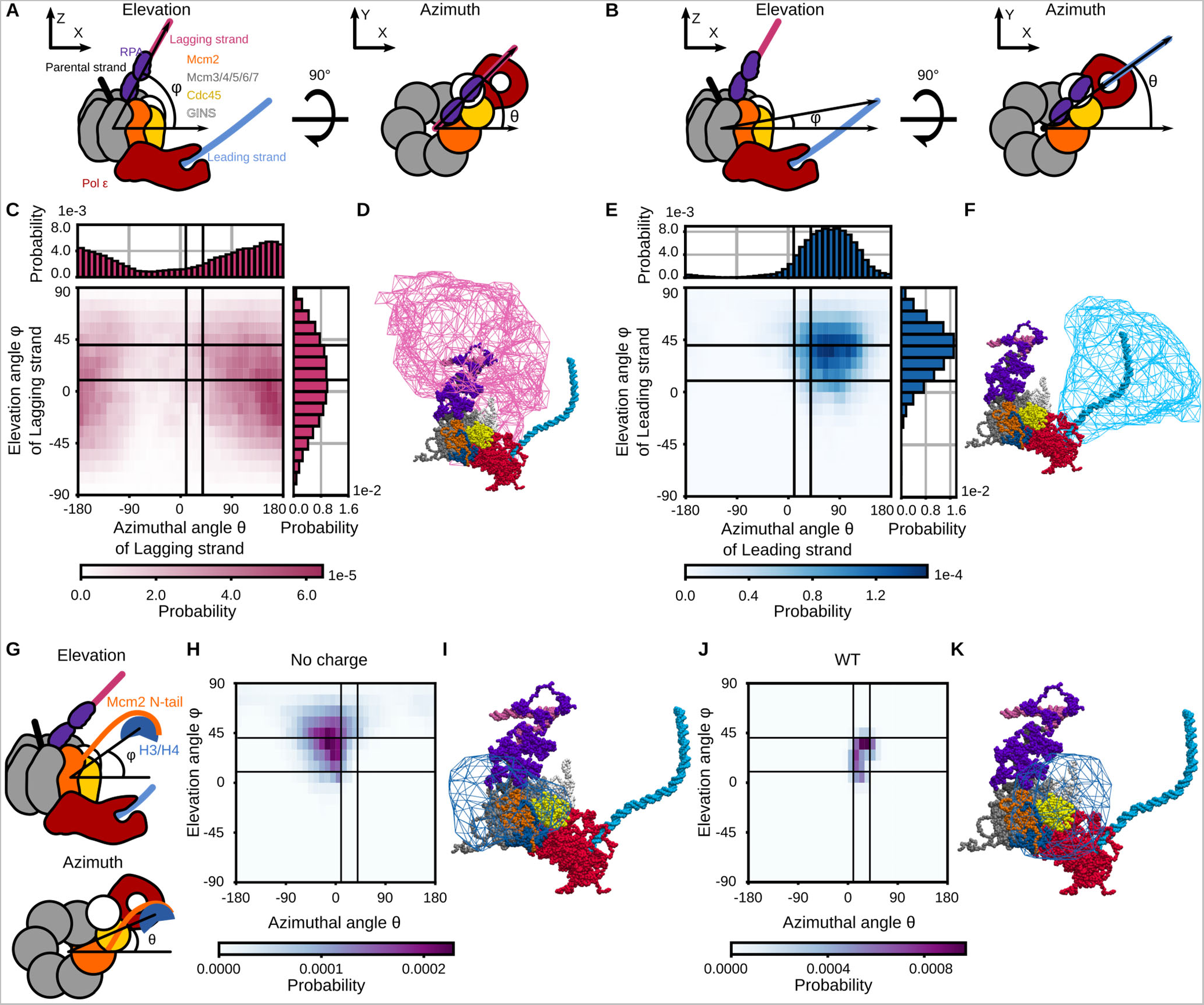
Distributions of orientations of the replicated strands and the H3/H4 tetramer in the simulations of the replicated-DNA-engaged replisome. **(A, B)** Schematic illustrations of the replicated-DNA-engaged replisome, which explains the definition of the elevation (*φ*) and azimuthal (*θ*) angles to describe the lagging (A) and leading (B) strand orientation. The color scheme is the same as in Figure 1B. **(C, E)** 1D and 2D probability distributions of the lagging (C) and leading (E) strand orientation calculated from the simulation trajectories until recycling. **(D, F)** Iso-surfaces of the spatial probability density of the lagging (D) and leading (F) strand orientation. The iso-value was set to 0.0001 [Å^−3^]. **(G)** Schematic illustrations of the replicated-DNA-engaged replisome explaining the definition of the elevation (*φ*) and azimuthal (*θ*) angles to describe the H3/H4 tetramer orientation. **(H, J)** 2D distributions of the H3/H4 tetramer orientation calculated from trajectories of the simulations in the absence (H) and presence (J) of charges in the Cdc45 acidic loop. **(I, K)** Iso-surfaces of spatial probability distributions of the H3/H4 tetramer orientation in the simulation in the absence (I) and presence (K) of charges in the Cdc45 acidic loop. The iso-value was set to 0.0001 [Å^−3^]. **(C, E, H, J)** The black auxiliary lines, which encircle the populated area in (J), mark the azimuthal angles of 10° and 40° and the elevation angles of 10° and 40°.

Next, we performed a similar analysis using the vector from the COM of Mcm2-7 to the H3/H4 tetramer to calculate the elevation (φ) and azimuthal (θ) angles (Figure 3G). When the H3/H4 tetramer did not associate with Cdc45, the vector tended to orient to the direction with an elevation angle of 43° ± 20° and an azimuthal angle of −11° ± 48° (Figure 3H & I). On the other hand, when the H3/H4 tetramer was associated with Cdc45, the vector tended to orient to the direction with an elevation angle of 25° ± 13° and an azimuthal angle of 25° ± 12° (Figure 3J & K). Therefore, the elevation angle shifted from 43° ± 20° to 25° ± 13° upon Cdc45 association while the azimuthal angle shifted from −11° ± 48° to 25° ± 12°. The region where the DNA strands (Figures 3D & 3F) or the H3/H4 tetramers (Figures 3I and 3K) frequently resided was surrounded by a polygon surface. In the simulations with a charged Cdc45 acidic loop (WT), the region where the H3/H4 tetramers resided (Figure 3J and 3K) has an about five times larger overlap with the region of the leading strand (Figure 3E and 3F, 0.70 %) than that of the lagging strand (Figure 3C and 3D, 0.14 %). Thus, the increase in co-orientation probability contributes to the leading strand bias of histone recycling in the Cdc45-mediated pathway.

### The number of RPA on the lagging strand regulates the strand bias

Previous studies have shown that the number of RPA on the lagging strand inversely correlates with the concentration of Pol α^54,62^. Although a single-molecule fluorescence imaging study revealed that the number of RPA is 1.5 ± 0.3 at 70 nM Pol α^54^, it may fluctuate in a cellular environment. To investigate the dependency of the number of RPA on recycling, we performed molecular dynamics simulations for 1 × 10^8^ steps using the new sets of initial structures: one without RPA (0-RPA, 16 nt ssDNA gap on lagging strand) and one with a single molecule of RPA (1-RPA, 36 nt ssDNA gap) (Figure 4A & B). As a result, the Mcm2 N-tail deposited the H3/H4 tetramer on the leading or lagging strand via the Cdc45-mediated or unmediated pathway (Figure 4C & Supplementary Figure 5A–5D) regardless of the number of RPA.

**Figure 4:**
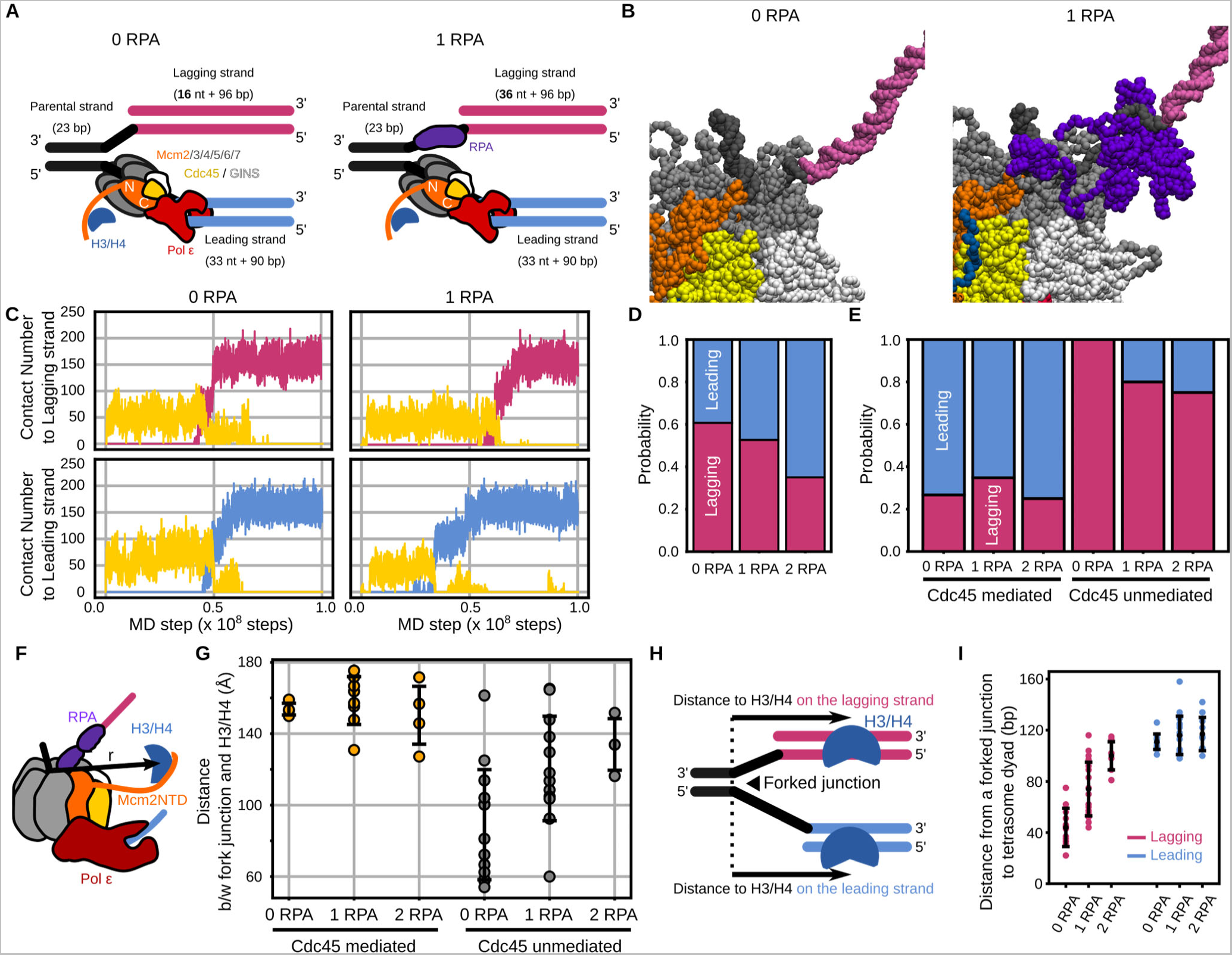
The H3/H4 tetramer was deposited on the replicated strands by the Mcm2 N-tail in the simulations of the replicated-DNA-engaged replisome with zero or one RPA. **(A)** Schematic illustration of the initial structures. The color scheme is the same as in Figure 1B. **(B)** Initial structures with zero (left) and one (right) RPA molecule. **(C)** The number of residues in the H3/H4 tetramer contacting the lagging strand (pink), the leading strand (cyan), and Cdc45 (yellow) in the case of zero (left) and one (right) RPA molecule. **(D)** Ratios of the replicated strands on which the H3/H4 tetramer was deposited in the simulations with zero, one, and two RPA molecules. **(E)** The same as in (D), but trajectories were classified into the Cdc45-mediated and unmediated pathways. **(F)** Schematic illustration explaining the definition of the 3D distance from the fork junction to the H3/H4 tetramer. **(G)** Distances from the fork junction to the H3/H4 tetramer at the moment of recycling to the lagging strand via the Cdc45-mediated pathway (orange) and the Cdc45-unmediated pathway (gray). **(H)** Schematic illustration explaining the definition of the distance from the fork junction to the destination location. **(I)** Distances from the fork junction to the position of the tetrasome dyad on the lagging (pink) and leading strand (cyan) in the simulations with zero, one, and two RPA molecules.

In the 0-RPA case, 56% (28/50) of the trajectories showed recycling to the leading or lagging strands, while 2% (1/50) and 42% (21/50) resulted in binding to the parental strand and no binding, respectively. Of the 28 recycling trajectories, 54% (15/28) and 46% (13/28) showed recycling via the Cdc45-mediated and unmediated pathways, respectively. 100% (13/13) of the recycling trajectories via the Cdc45-unmediated pathway resulted in the deposition of the H3/H4 tetramer on the lagging strand, while 73% (11/15) of the trajectories via the Cdc45-mediated pathway ended up with the deposition on the leading strand.

In the 1-RPA case, 38% (38/100) of the trajectories showed recycling to the leading or lagging strands, while 23% (23/100) and 39% (39/100) resulted in binding to the parental strand and no binding, respectively. Of the 38 recycling trajectories, 61% (23/38) and 39% (15/38) showed recycling via the Cdc45-mediated and unmediated pathways, respectively. 80% (12/15) of the trajectories of recycling via the Cdc45-unmediated pathway resulted in the deposition of the H3/H4 tetramer on the lagging strand, while 65% (15/23) of the trajectories via the Cdc45-mediated pathway ended up with the deposition on the leading strand.

These statistics showed that the lagging-strand recycling bias was mitigated as the number of RPA associated with the lagging strand increased (Figure 4D). To get further insights, we classified the trajectories based on the pathways. Note that the H3/H4 tetramer was deposited mainly on leading and lagging strands via the Cdc45-mediated and unmediated pathways, respectively, as in the 2-RPA case. The analysis showed that the leading strand bias via the Cdc45-mediated pathway was unaffected, while the lagging strand bias via the Cdc45-unmediated pathway was mitigated (Figure 4E). Together, the simulations suggested that RPA association with a lagging strand promotes the leading strand bias by inhibiting recycling to the lagging strand via the Cdc45-unmediated pathway.

To confirm whether RPA binding affects the dynamics of the H3/H4 tetramer, we analyzed the orientation of the H3/H4 tetramer until recycling in the 0- and 1-RPA case as in the 2-RPA case above. The H3/H4 tetramer which did not bind to Cdc45 oriented with the azimuthal angle of −13° ± 44°, −18° ± 50°, and −18° ± 34° and with the elevation angle of 35° ± 23°, 39° ± 22°, and 39° ± 19° in 0-, 1-, and 2-RPA case (Supplementary Figure S6A-C). On the other hand, the H3/H4 tetramer which bound to Cdc45 oriented with the azimuthal angle of 20° ± 11°, 7° ± 12°, and 25° ± 13° and with the elevation angle of 9° ± 13°, 19° ± 13°, and 25° ± 12° in the 0-, 1-, and 2-RPA case (Supplementary Figure S6D-F). We further calculated the three-dimensional distance between the H3/H4 tetramer and the fork junction until recycling (Figure 4F). The distances were 133 ± 32 Å, 129 ± 35 Å, and 137 ± 25 Å in the 0-, 1-, and 2-RPA case, respectively, when the H3/H4 tetramer dissociated from Cdc45 (Supplementary Figure S6G-I). On the other hand, the distances were 150 ± 14 Å, 150 ± 15 Å, and 150 ± 15 Å, whereas the H3/H4 tetramer associates to Cdc45. Therefore, these analyses indicated that RPA binding did not significantly affect the region where the H3/H4 tetramer migrates until recycling. We performed a similar analysis on the dynamics of lagging strands and found that the direction of lagging strands is also unaffected by RPA binding (Supplementary Figure S7A-B & Figure 3C).

To investigate the reason for the bias mitigation, we analyzed the distance between the fork junction and the H3/H4 tetramer at the moment of recycling to the lagging strand. (Figure 4F). The distance was 89 ± 31 Å, 121 ± 29 Å, and 134 ± 14 Å via the Cdc45-unmediated pathway, while 154 ± 3 Å, 159 ± 13 Å, and 150 ± 16 Å in the 0-, 1-, and 2-RPA case (Figure 4G). In other words, in the absence of RPA, the distance was widely distributed from 60 Å to 160 Å, whereas as the number of RPA molecules increased, the average value increased, and the range became narrower, which may be caused by occlusion and extension of ssDNA regions by RPA (Supplementary Figure S6J-K). These results suggested that the migration area of the histones that can be recycled is constrained as the number of RPA molecules increases. It is reasonable to think that this constraint is one reason RPA prevents recycling to the lagging strand via the Cdc45 unmediated pathway.

Next, we focused on the destination location where the H3/H4 tetramer was recycled. We defined the distance from the fork junction as the number of nucleotides from the fork junction to the dyad of a recycled tetrasome (Figure 4H). As the number of RPA associated with the lagging strand increased from zero to two, the distance increased from 43 ± 15 to 100 ± 11 nt for recycling to the lagging strand (Figure 4I). On the other hand, the distance was unaltered in a significant way for recycling to the leading strand. This is as expected since the binding of RPA to the lagging strand shifts the naked dsDNA region required for recycling upstream (Figure 1B & 4A). Remarkably, simulation results revealed that the H3/H4 tetramer could be recycled to the lagging strand even in the presence of two RPA molecules, equal to or slightly more than the average number (1.5 ± 0.3) in physiological conditions^54,55^.

### Pol ε association affects the destination location for the leading strand recycling

In the replicated-DNA-engaged replisome structure^27,28^, Pol ε attaches to the CMG helicase complex and bends the leading strand (Figure 1D & 5A). To investigate the role of DNA bending in recycling, we performed 50 runs of simulations of the replicated-DNA-engaged replisome without Pol ε and RPA molecules for 1 × 10^8^ steps (Figure 5A). As expected, Pol ε restrained the leading strand orientation in the simulations (Figure 5B & 5C), and the leading strand oriented to the direction with an elevation angle of 34° ± 25° and an azimuthal angle of 71° ± 49° in the presence of Pol ε (CMGE) while with an elevation angle of −24° ± 33° and an azimuthal angle of −178° ± 95° in its absence (CMG).

**Figure 5:**
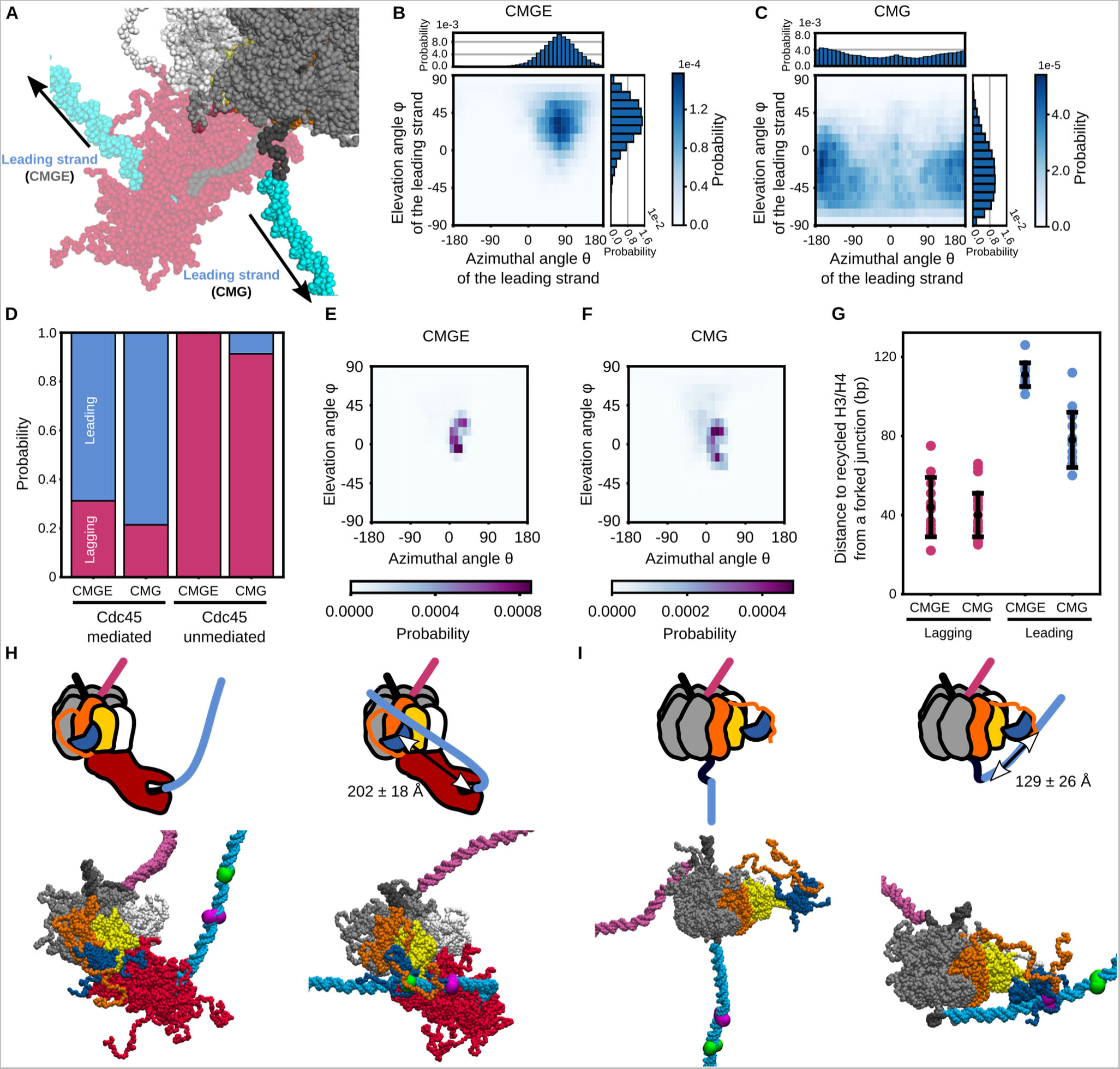
The H3/H4 tetramer was deposited on the daughter strands by the Mcm2 N-tail in the simulations of the replicated-DNA-engaged replisome without Pol ε and RPA. **(A)** The structure of CMG (opaque) superimposed to that of CMGE (transparent). **(B, C)** 1D and 2D probability distributions of the leading strand orientation calculated from the simulations of CMGE and CMG. **(D)** Ratios of the replicated strands on which the H3/H4 tetramer was deposited in the simulations of CMGE and CMG. **(E, F)** 2D distributions of the H3/H4 tetramer orientation calculated from trajectories of the simulations of CMGE and CMG **(G)** Distances from the fork junction to the deposited positions on the lagging (pink) and leading strand (cyan) in the simulations of CMGE and CMG. **(H, I)** Schematic illustrations and representative structures of CMGE (H) and CMG (I). The initial structure and the structure at the exact moment of the H3/H4 deposition are shown on the left and right, respectively. The color scheme is the same as in Figure 1B & 1D. Additionally, the green and magenta spheres on the leading strand represent the average recycled position in the CMGE and CMG simulations, respectively.

In the simulations without Pol ε, 74% (37/50) of the trajectories showed recycling to the leading or lagging strands, while 6% (3/50) and 20% (10/50) resulted in binding to the parental strand and no binding, respectively. Of the 37 recycling trajectories, 38% (14/37) and 62% (23/37) showed recycling via the Cdc45-mediated and the Cdc45-unmediated pathways, respectively. The H3/H4 tetramer preferred to be recycled via the Cdc45-unmediated pathway in the absence of Pol ε, while it preferred to be recycled via the Cdc45-mediated pathway in the presence of Pol ε. 79% (11/14) of the trajectories of recycling via the Cdc45-mediated pathway resulted in the deposition of the H3/H4 tetramer on the leading strand, while 91% (21/23) of the trajectories via the Cdc45-unmediated pathway ended up with the deposition on the lagging strand (Figure 5D). These statistics are comparable to those from the simulations in the presence of Pol ε (69% and 100%, respectively, Figure 5D). Together, DNA bending by Pol ε did not significantly affect the strand bias.

Next, we performed the analysis using the vector from the COM of Mcm2-7 to the H3/H4 tetramer to calculate the elevation (φ) and azimuthal (θ) angles to investigate the effect of Pol ε on the movement of the H3/H4 tetramer. The vector tended to orient to the direction with an elevation angle of 15° ± 19° and an azimuthal angle of 15° ± 26° in the presence of Pol ε (Figure 5E). On the other hand, the vector tended to orient to the direction with an elevation angle of 17° ± 27° and an azimuthal angle of 12° ± 32° in the absence of Pol ε (Figure 5F). Therefore, Pol ε caused a minuscule change in the orientation of the H3/H4 tetramer.

To investigate the destination location in the absence of Pol ε, we calculated the distance from the fork junction to the recycled position as defined above (Figure 4H). The distances were 78 ± 13 and 40 ± 11 nt for the leading and lagging strands, respectively (Figure 5G). Thus, the distance for the leading strand in the absence of Pol ε decreased compared to its presence while not significantly altered for the lagging strand. Since we did not change the leading ssDNA length regardless of Pol ε existence, and the Pol ε only occludes the ssDNA region, its occlusion alone cannot explain the alteration of the destination location. Instead, the Pol ε binding extended the ssDNA region and moved the ssDNA-dsDNA junction away from the H3/H4 tetramer in the Cdc45-mediated pathway, where the tetramer is preferably recycled to the leading strand (Figure 5H & 5I). In fact, we measured the distance between the junction and Cdc45, which turned out to be 202 ± 18 Å and 129 ± 26 Å in the presence and the absence of Pol ε. This result indicated that the temporal dissociation of Pol ε, e.g., upon polymerase exchange^54^, may alter the destination location and, hence, the gene regulation in daughter cells.

We further simplified the system by excluding Cdc45, GINS, Pol ε, and RPA to investigate the effect of leading strand orientation on recycling. This system, which only contains the replicated DNA and Mcm2-7, allowed us to simulate various conditions. First, the ends of the leading and lagging strands were fixed in space with a harmonic potential (k = 0.002 kcal/mol·Å) so that the leading strand orients with an elevation angle of −90° and the lagging strand orients with varying angles (φ = −45°–75° with 30° steps, θ = −135°–180° with 45° steps) (Figure 6A). From each of these 40 initial structures, we repeated ten runs of simulations for 1 × 10^8^ steps. Almost all the trajectories showed recycling to the lagging strand (189/400), association with the parental strand (157/400), or no association (53/400). (Figure 6B & 6C). On the other hand, the recycling probability to the leading strand is vanishingly low (1/400) regardless of the lagging strand orientation (Figure 6B). Thus, the deviation from the seemingly natural direction of the leading strand is essential for recycling to the leading strand. Next, the ends of the leading and lagging strands were fixed so that the lagging strand orients with an elevation angle of 90° and the leading strand orients with varying angles (φ = −75°–45° with 30° steps, θ = −135°–180° with 45° steps) (Figure 6D). Again, from each of these 40 initial structures, we performed ten simulation runs for 1 × 10^8^ steps and observed recycling to the leading strand in 13% trajectories (Figure 6E & 6F). Interestingly, the recycling probability to the leading strand increased as the elevation angle of the DNA became larger, and the azimuthal angle was closer to zero (Figure 6E). However, by comparing Figure 5B and 6E, we can see that Pol ε fixes the leading strand orientation to the suboptimal direction for recycling, which explains the finding that DNA bending by Pol ε does not significantly alter the strand bias of recycling in the Cdc45-unmediated pathway.

**Figure 6:**
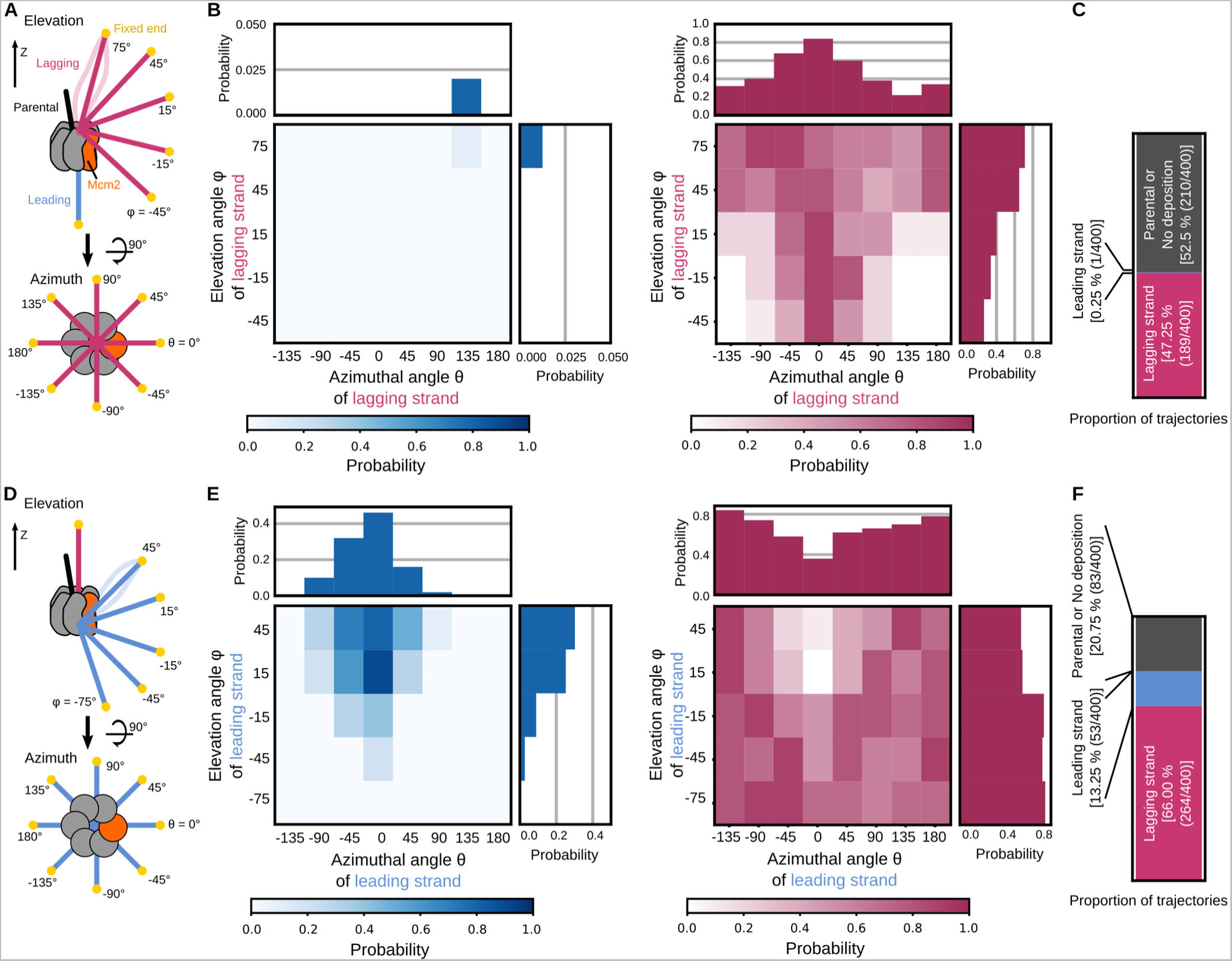
The H3/H4 tetramer was deposited on the daughter strands by the Mcm2 N-tail in the simulations of the replicated-DNA-engaged replisome without Cdc45, GINS, Pol ε, RPA and with the leading or lagging strand orientation loosely constrained in space. **(A)** Schematic illustrations of the initial structures in which the orientation of the lagging strands was loosely constrained. The color scheme is the same as in Figure 1B. **(B)** 1D and 2D distributions of the probability of the leading (left) and lagging (right) strand histone deposition in the simulations with the varying lagging strand orientation. **(C)** Ratios of the strands to which the H3/H4 tetramer was recycled in the simulations with the varying lagging strand orientations. **(D)** Schematic illustrations of the initial structures in which the orientation of the leading strands was loosely constrained. The color scheme is the same as in Figure 1B. **(E)** 1D and 2D distributions of the probability of the leading (left) and lagging (right) strand histone deposition in the simulations with the varying leading strand orientation. **(F)** Ratios of the strands to which the H3/H4 tetramer was recycled in the simulations with the varying leading strand orientations. The color scheme is the same as in Figure 1B.

## DISCUSSION

In this study, we performed molecular dynamics simulations of a yeast DNA replication machinery containing Mcm2-7, Cdc45, GINS, Pol ε, and RPA, bound to an H3/H4 tetramer and replicated DNA to visualize the structural trajectory from the H3/H4 tetramer bound to Mcm2 until recycled to the replicated strands. The simulations and the *in vitro* replication assays combinatorially showed that Mcm2 can directly deposit the H3/H4 tetramer onto the replicated strands without additional factors such as histone chaperones. Interestingly, the simulations also suggested that the H3/H4 tetramer is more likely recycled to the leading and lagging strands in the Cdc45-mediated and unmediated pathways, respectively. Also, RPA binding to the lagging strand inhibited recycling via the Cdc45-unmediated pathway, tilting the strand bias toward the leading strand. On the other hand, Pol ε binding to the leading strand did not significantly alter the strand bias but did affect the destination location on the leading strand.

The most prominent prediction from the simulations is the Cdc45-mediated pathway of recycling. In the simulations, the H3/H4 tetramer was associated with the acidic loop of Cdc45 and was mainly recycled to the leading strand. The Native-PAGE assays supported that Cdc45 and an H3/H4 tetramer interact electrostatically. The interaction may have a regulatory role in the strand bias. Remarkably, the T189 and T195 residues in the acidic loop of Cdc45 are known to be phosphorylated, a prerequisite for recruiting Rad53, an S-phase checkpoint kinase^61^. These post-translational modifications lead to more negative charges in the acidic loop, which may enhance interactions with the H3/H4 tetramer. Also, the binding of Rad53 to the acidic loop may weaken the binding of the H3/H4 tetramer. The regulations of the strand bias of histone recycling by post-translational modifications and protein binding are a new paradigm of epigenetic inheritance, and experimental verification is strongly desired.

The previous *in vitro* replication assays using DNA substrates with multiple nucleosomes revealed that histone chaperone FACT and chromatin remodeler INO80 and ISW1 are required for efficient chromatin replication^19^. On the other hand, the *in vitro* replication assays in our current study showed that a replisome progresses on one nucleosome without histone chaperones and chromatin remodelers. These results collectively suggested that each nucleosome acts as a fragile barrier against replication and that the effects are additive. In some simulation trajectories, the H3/H4 tetramer was stably deposited onto the parental strand, which may act as a fragile barrier. Weakening this binding may be the role that FACT, INO80, and ISW1 played for efficient chromatin replication in the previous *in vitro* replication assays^19^.

Previous studies revealed that various components of a replisome and histone chaperones regulate the histone recycling efficiency and the strand bias^14,18,21,22,63–65^. Of them, the SCAR-seq studies demonstrated that Mcm2 contributes to recycling, especially on the lagging strand, in the cellular condition^14,22^. The current study showed that the Cdc45-mediated and unmediated pathways preferably deposit the H3/H4 tetramer on the leading and lagging strands, respectively. These results collectively suggested that the predominant pathway in the cellular condition is the Cdc45-unmediated pathway. As a limitation of this study, the simulation system lacks the replisome components such as Ctf4, Pol α, Fen1, Lig1, and PCNA and the histone chaperones such as CAF-1, Asf1, FACT, and HJURP, which were suggested to cooperate with Mcm2 for histone recycling^18,22,26,63–66^. It is tempting to assume that these additional factors are decisive in choosing the dominant pathway.

In the simulations with Pol ε, the H3/H4 tetramer was deposited on the leading strand at the position 111 nt away from the fork junction. In our structural model, the length of the ssDNA gap on the leading strand was 33 nt and passed through the central channel of Mcm2-7 and the catalytic subunit of Pol ε. In this configuration, the ssDNA region that emerges from Mcm2-7 and enters Pol ε was the shortest length required and stretched, but it is possible that this region becomes longer in the cellular condition. A recent study has shown that this ssDNA region is elongated due to uncoupling between DNA unwinding and leading strand synthesis upon replication stress^53^. This ssDNA looping between Mcm2-7 and Pol ε may make the destination location further away from the fork junction. This consideration leads to an attractive hypothesis that the replication speed modulates the transcriptional programs in daughter cells via altered nucleosome positioning.

RPA binding to the ssDNA region on the lagging strand hindered recycling to the lagging strand via the Cdc45-unmediated pathway. Also, this binding caused the H3/H4 tetramers to be recycled to positions on the lagging strand further away from the fork junction. Previous *in vitro* and *in vivo* studies have shown that the ssDNA length on the lagging strand varies between 0 to 2,000 nt depending on the concentration of Pol α-primase complex^54,62^, which is required to initiate the lagging strand synthesis. This knowledge and our simulation results collectively suggested that the lagging strand synthesis initiation rate modulates the strand bias. Notably, a recent ChIP-NChAP study has shown that the parental histones preferentially bind to the strand replicated first^67^ in line with this suggestion.

In this study, we performed molecular dynamics simulations of a replisome already bound to an H3/H4 tetramer. Since the previous studies showed that the Mcm2 N-terminal tail binds to an H3/H4 tetramer^15,16^ and contributes to the recycling of a parental H3/H4 tetramer, especially to the lagging strand, it is reasonable to assume that the bound structure is the genuine intermediate state of histone recycling in the cellular condition^14,22^. However, we cannot rule out the possibility that the newly synthesized histones also bind to the Mcm2 N-tail. The simulations suggested that even newly synthesized histones can be deposited on the replicated strands if attached to the Mcm2 N-tail. Whether the binding of the H3/H4 tetramer is limited only to the parental one, and if so, what is the molecular mechanism to accomplish it remain intriguing open questions. Notably, previous single-molecule imaging using *Xenopus laevis* egg extracts revealed that the recycling efficiency of the parental histones depends on the concentration of the newly synthesized histones^20^, supporting that the pathways simulated in this study are used to deposit both parental and newly synthesized histones.

In recycling, either of the daughter strands took the H3/H4 tetramer away from the Mcm2 N-tail. Thus, the extent to which the strands probe the H3/H4 tetramer around a replication fork is critical. A recent study has proposed a theoretical model that diffusion of parental histones or DNA segments accounts for dispersed histone inheritance in active chromatin domains^68^. In this diffusion-driven model, histones can be recycled on any physically proximal DNA segment. Notably, our simulations are in line with the possibility that DNA segments far away from the fork junction associate with H3/H4 tetramer bound to the Mcm2 N-tail. Also, a previous *in vitro* single-molecule imaging suggested that histones can be recycled on the destination location further than the DNA persistent length via DNA loop formation^69^. Therefore, the pathways visualized in the current simulations may contribute to the dispersion of epigenetic marks in active chromatin domains.

The coarse-grained model used in this study allowed us to simulate the dynamics of a large molecular system like a replisome for a reasonable timescale to observe the H3/H4 recycling to the replicated strands. Although previous and current studies have carefully calibrated the potential energy functions and their parameters for the intra- and inter-molecular interactions, the treatment of the inter-molecular interactions is relatively simple. This simple treatment may underestimate the moderately strong interaction between the H3/H4 tetramer and RPA, which was experimentally detected, for example^70^. On this occasion, additional structural information may help improve the accuracy of the simulation results. Otherwise, the potential energy functions for hydrogen bonding and hydrophobic interactions can be incorporated^33,71^. Also, the simulations cannot account for the coupling among DNA unwinding, DNA synthesis, and histone recycling. Future studies can address this limitation by incorporating the potential switching procedure to model protein conformational change upon ATP hydrolysis^72^. Despite these limitations, the simulations provided insights and experimentally testable predictions on the molecular mechanism, which is vital for elucidating the intracellular histone recycling mechanisms regulated by the cooperation of various histone chaperones.

## MATERIALS AND METHODS

### Coarse-grained molecular dynamics simulations

For proteins (Cdc45, Mcm2-7, GINS, Pol ε, RPA, and an H3/H4 tetramer), we used the AICG2+ model^35^ representing one amino acid as one bead located at C_α_ atom position. The following paragraphs describe how we modeled the initial structures of the CMG helicase complex (Cdc45 + Mcm2-7 + GINS), DNA polymerase ε (Pol ε), Replication protein A (RPA), and the H3/H4 tetramer.

To prepare the initial structure of the CMG helicase complex, we used the cryo-EM structure (PDB ID: 6U0M)^27^ as a reference. The CMG helicase comprises three proteins: Cdc45, Mcm2-7, and GINS^40^ (Figure 1A). The reference structure contains the residues 1–650 of Cdc45. Mcm2-7 is a protein consisting of six subunits: Mcm2/3/4/5/6/7. The reference structure contains the residues 1–868 of Mcm2, 17–738 of Mcm3, 177–838 of Mcm4, 24–693 of Mcm5, 103–840 of Mcm6, and 1–729 of Mcm7. GINS is a protein consisting of four subunits: Sld5, Psf1, Psf2, and Psf3. The reference structure contains the residues 3– 294 of Sld5, 1–208 of Psf1, 3–200 of Psf2, and 3–193 of Psf3. We treated the residues 166-217 and 437– 457 of Cdc45, 1–200 and 707-736 of Mcm2, 58–90, 142–150, 332–337 and 571–650 of Mcm3, 213–220, 470–497, 731–740, 780–792 and 839–850 of Mcm4, 104–129, 212–234, 306–318, 340–345 and 644–646 of Mcm5, 246–259, 415–427, and 484–509 of Mcm6, 32–58, 159–188, 217–219 and 387–392 of Mcm7, 3–53, 111–120 and 239–247 of Sld5, 33–49 of Psf2, and 30–32, 59–67 and 142–161 of Psf3 as intrinsically disordered regions. The initial conformations of the intrinsically disordered regions (IDRs) were generated using MODELLER^73^. All IDRs in other protein models were similarly generated.

For Pol ε, we used the cryo-EM structure (PDB ID: 6HV9^28^) as a reference. Pol ε comprises four proteins: Pol2, Dpb2, Dpb3, and Dpb4^41^ (Figure 1A). The reference structure contains the residues 1308– 2222 of Pol2 and 1–90 of Dpb2 interacting with the CMG helicase complex. To obtain the full-length Pol ε reference structure containing the residues 1–2222 of Pol2, 1–689 of Dpb2, 1–201 of Dpb3, and 1–196 of Dpb4, we superimposed the cryo-EM structure of Pol ε holoenzyme (PDB ID: 6WJV^41^) and Dpb2 (PDB ID: 6HV9^28^) to the reference structure above (PDB ID: 6U0M^27^). We treated the residues 1–30, 91– 107, 215–233, 664–677, 1187–1269, 1393–1403, 1748–1783, 1977–1993, 2033–2042, 2073–2099, and 2122–2127 of Pol2, 91–169, 195–206, 234–265, 368–377 and 557–597 of Dpb2, 1–8 and 94–201 of Dpb3, 1–17 and 125–196 of Dpb4 as IDRs.

For the H3/H4 tetramer, we used the crystal structure of a nucleosome core particle (PDB ID: 1KX5^49^) as a reference. We treated the residues 1–32 of H3 and 1–23 of H4 as IDRs.

For RPA, the partial crystal, cryo-EM, and homology-modeled structures were connected by flexible linker regions with MODELLER^73^. RPA is a protein consisting of three subunits: Rfa1, Rfa2, and Rfa3 (Figure 1A). The reference structure of Rfa1 comprises the partial structures of DBD-F (residues 1–132, PDB ID: 5OMB^44^), DBD-A (residues 181–294, PDB ID: 1YNX^45^), DBD-B (residues 295–419, PDB ID: 1JMC^46^, homology model), and DBD-C (residues 442–619, PDB ID: 6I52^47^). The reference structure of Rfa2 comprises the partial structures of the winged helix domain (residues 205–273, PDB ID: 4OU0^48^, homology model) and DBD-D (residues 32–182, PDB ID: 6I52^47^). The reference structure of Rfa3 comprises the structure of DBD-E (residues 1–122, PDB ID: 6I52^47^). The structures of Rfa1, Rfa2, and Rfa3 were assembled by superimposing DBD-C of Rfa1, DBD-D of Rfa2, and DBD-E of Rfa3 to the heterotrimeric cryo-EM structure (PDB ID: 6I52^47^). We treated the residues 133–180 and 420–441 of Rfa1, 1–31, and 183–204 of Rfa2 as IDRs.

For DNA, we used the 3SPN.2 model^36^, in which one nucleotide was represented as three beads located at the centroid of base, sugar, and phosphate groups. The replicated-DNA structure was modeled by superimposing the ideal B-form DNA structure generated using 3DNA^74^ to the forked DNA in the cryo-EM structure (PDB ID: 6U0M^27^), which comprises 23 bp, 90 bp, and 96 bp of dsDNA for the parental, leading, and lagging strands and 33 nt and varying length (16 nt in 0-RPA, 36 nt in 1-RPA, and 56 nt in 2-RPA) of ssDNA for the leading and lagging strands. The structure-based potential was applied to stabilize the B-form DNA structure and to reproduce the persistence length of ds and ssDNA, the melting temperature, and the hybridization rate.

The potentials for the excluded volume and electrostatic interactions were applied to the inter-molecular interactions. On top of them, the structure-based potential was applied to the protein residue pairs or the residue-nucleotide pairs which form contact in the experimentally solved structures to stabilize the cryo-EM structure of the RPA-ssDNA complex (PDB ID: 6I52^47^), the crystal structure of the Pol ε-DNA complex (PDB ID: 4M8O^75^), and the crystal structure of the H3/H4 dimer-Mcm2 complex (PDB ID: 4UUZ^15^). The potential for the hydrogen bonding interactions was also applied to the interactions between the H3/H4 tetramer and DNA. The parameters of this potential were calibrated in the previous studies^33,34^ to stabilize the canonical nucleosome structure and to reproduce the salt-concentration-dependent DNA unwrapping from a nucleosome.

The potential derived from Debye-Hückel’s theory was applied to the electrostatic interactions. We arranged the partial charges on protein surface beads by the RESPAC algorithm^32^ so that the model reproduced the electrostatic potential around the all-atom structures except for the beads in the disordered regions where we set +1*e* charge on lysine and arginine residues, −1*e* charge on aspartic acid and glutamic acid residues, and zero charge on the other residues. The charges of DNA phosphate groups were set to −0.6*e* for intra-DNA interactions to model the counter ion condensation around the phosphate groups within the framework of the Debye-Hückel model. On the other hand, the phosphate charges were set to −1.0*e* for protein-DNA interactions to account for releases of counter ions upon the association between DNA and a protein.

We performed Langevin dynamics simulations to integrate the equations of motion with a timestep of 0.3 CafeMol time units (~14.7 fs). Temperature and the friction constant were set to 300K and 0.843, respectively. The monovalent ion concentration of the Debye-Hückel model was set to 300 mM, and the dielectric constant was set to 78.0. All the simulations were conducted using CafeMol3.2^56^ (https://www.cafemol.org). It took ~2 weeks to compute the entire system for 1 × 10^8^ steps using two CPU cores (Intel^®^ Xeon^®^ Gold 6326) in parallel.

### Optimization of the potential for the interactions between Mcm2 and the H3/H4 tetramer

In addition to the potentials for the excluded volume and electrostatic interactions, the AICG2+ potential was applied for the inter-protein interactions between the Mcm2 histone-binding region (residues 69-121) and the H3/H4 tetramer based on the crystal structure (PDB ID: 4UUZ^15^). The non-local term in the AICG2+ potential function is given as

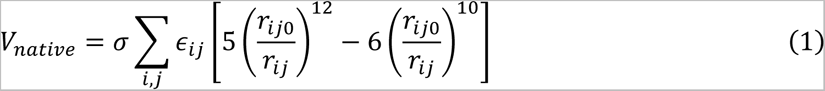

where σ is a global scaling parameter, *ε_ij_* is a site-specific parameter between the *i*-th residue in Mcm2 and the *j*-th residue in an H3/H4 dimer, and *r_ij0_* and r*_ij_* are distances between the *i*-th and the *j*-th residue in the native and the simulated structure, respectively. Since this interaction plays a decisive role in recycling, we sought to carefully calibrate the scaling parameter σ so that the experimentally measured dissociation constant is reproduced. Thus, we performed replica-exchange molecular dynamics simulations of the Mcm2 N-tail (residues 1–200) and the H3/H4 dimer in a sphere with a radius of 100Å for 1 × 10^9^ steps with σ set to 0.65, 0.70, and 0.75 (Supplementary Figure 1A & 1B). The temperature of each of the ten replicas was set from 300K to 390K with a 10K linear increment. To define the bound and unbound states, we calculated the *Q*-score of each simulation snapshot (Supplementary Figure 1C). The *Q*-score was defined as the ratio of the number of residue-residue contacts to the number of natively formed contacts^15^. We considered that a residue pair forms a contact when these beads are within 1.2 times the distance in the native structure. Then, we defined the bound state as the state with *Q* > 0. By comparing the binding free energy calculated from each simulation (ΔG*^σ=0.65^* = −5.3 ± 0.1 kcal/mol, ΔG*^σ=0.70^* = −10.0 ± 0.3 kcal/mol, and ΔG*^σ=0.75^*= −15.7 ± 0.5 kcal/mol) and experimental measurement (ΔG*^expt^* = −10.4 kcal/mol), we decided to set σ to 0.70, which best reproduced the experiment.

### Simulations in which orientation of the leading or lagging strand was constrained

We performed the simulations in which the leading or lagging strand was constrained to be oriented in a variety of directions. First, we moved the molecules so that the COM of Mcm2-7 was on the origin and the longitudinal axis aligned to the *z*-axis. The COM of each subunit in Mcm2-7 was fixed in space by a harmonic potential (*k* = 0.1 kcal/mol·Å^2^). The ends of the leading and lagging strands were also loosely fixed in space with a harmonic potential (*k* = 0.002 kcal/mol·Å^2^) so that the leading strand orients with an elevation angle of −90° and the lagging strand orients with varying angles (*φ* = −45°–70° with 30° steps, *θ* = −135°–180° with 45° steps) (Figure 6A) and that the lagging strand orients with an elevation angle of 90° and the leading strand orients with varying angles (*φ* = −75°–45° with 30° steps, *θ* = −135°–180° with 45° steps) (Figure 6D).

### Experimental material preparations

The ARS1-W601 DNA used for *in vitro* DNA replication was chemically synthesized and cloned into a plasmid pEX-A2J2 vector. The ARS1-W601 linear DNA fragments (1155 bp) were obtained by Polymerase Chain Reaction (PCR). The budding yeast histone octamers (H3, H4, H2A, and H2B) and replication proteins (ORC, Cdc6, Cdt1-Mcm2-7, DDK, S-CDK, Sld2, Sld3-Sld7, Dpb11, Cdc45, GINS, PCNA, RFC, Mcm10, RPA, Polymerase α (Pol α), Polymerase ε (Pol ε), Polymerase δ (Pol δ), Ctf4, Csm3-Tof1 and Mrc1) were purified as described previously^59^. Cdc45-7HIS, Cdc45-7HIS, (e) and histone H3/H4 complex used for native gel electrophoresis assays (Supplementary Figure S3A) was expressed and purified from *Escherichia coli* as also described previously^60,61,76^.

### Replication assays

The nucleosomes were initially assembled on the ARS1-W601 linear DNA (1155 bp) by salt gradient dialysis as described previously^60^. In brief, 27.7 nM DNA and 354 nM histone octamer were mixed and dialyzed against 20 mM Tris-HCl (pH 7.5), 2 mM EDTA, 2 mM DTT, and 0.08% NP40 with a linear NaCl gradient from 2 M to 0.2 M. This was further dialyzed against 25 mM HEPES-KOH (pH 7.5), 0.5 mM EDTA, 1 mM DTT, 0.04% NP40. The nucleosome substrate was then diluted 2-fold in Mcm buffer (25 mM HEPES-KOH (pH 7.5), 1 mM dithiothreitol (DTT), 5 mM ATP, 7.5 mM Mg(OAc)_2_, 5% glycerol, 0.01% (w/v) NP-40, 0.1 mg/ml BSA) containing 100 mM KOAc, 15 nM Orc, 60 nM Cdt1-Mcm2-7, and 30 nM Cdc6. After a 15-minute incubation at 30 °C, 50 nM DDK was added, and incubation was continued for 15 minutes. The reaction was then 4-fold diluted in replication buffer (25 mM HEPES-KOH (pH 7.5), 1 mM DTT, 5 mM ATP, 7.5 mM Mg(OAc)_2_, 5% glycerol, 0.01% (w/v) NP-40, 0.1 mg/ml BSA, 0.1 mM CTP, 0.1 mM GTP, 0.1 mM TTP, 40 µM dATP, 40 µM dCTP, 40 µM dGTP, 25 µM dTTP and 15 µM biotin-dUTP) containing 60 mM KOAc, and proteins in final concentration of Orc (3.75 nM), Cdt1-Mcm2-7 (15 nM), Cdc6 (7.5 nM), DDK (10 nM), S-CDK (5 nM), Sld2 (30 nM), Sld3-Sld7 (20 nM), Dpb11 (30 nM), Cdc45 (40 nM), GINS (12.5 nM), Tof1-Csm3 (20 nM), Mrc1 (10 nM), Ctf4 (20 nM), RFC (20 nM), PCNA (20 nM), RPA (100 nM), Pol α (5 nM), Pol ε (2 nM), Pol δ (10 nM), and Mcm10 (5 nM). The reaction mixture (100 µl) was incubated at 30°C for 20 minutes. Note that the final KOAc concentration was ~100 mM. The reaction was adjusted to 50 mM NaCl and 2 mM CaCl_2_, then further digested by MNase (1 U/ul) at 30°C for 30 minutes. The reaction was mixed with 4 µL of 0.5 M EDTA, 2 µL of 10% (w/v) SDS, and 1 µL of 20 mg/mL protease K, then incubated at 37 °C for 20 minutes. The sample was mixed with 1/6 volume of native DNA dye (15% (w/v) ficol, 10 mM HEPES-KOH (pH 7.5), 0.05% orange G) and applied to 7.5% polyacrylamide gel electrophoresis in EzRun TG buffer (ATTO) at room temperature for 65 minutes at 21 mA. After dipping in 1/2x TBE for 5 minutes, the DNAs were transferred to the Zeta-Probe membrane (Bio-Rad) using a wet-transfer blotter at 80 V for 50 minutes at 4 °C. The membrane was crosslinked using a UV illuminator and soaked in blocking solution (Cytiva) at room temperature for 20 minutes. The membrane was incubated with Dylight680-conjugated streptavidin in TBS containing 0.1% (w/v) Tween 20 and 0.01% (w/v) SDS for 30 minutes. After washing the membrane three times with TBS+Tween+SDS, gel images were captured by the ChemiDoc Touch imager (Bio-Rad).

### Native polyacrylamide gel electrophoresis assays

We mixed 0.5 μM Cdc45-7HIS and 0-2 μM H3/H4 tetramer in 20 μL reaction buffer (25 mM HEPES-KOH (pH7.5), 1 mM EDTA, 10% Glycerol, and 0.05% Tween 20) with 150, 300, and 750 mM KCl for low-, medium-, and high-salt conditions, respectively. The reactions were incubated for 15 minutes at 30 ℃ and divided into two 10 μL aliquots. Each aliquot was run on 5–20% Tris-Glycine or sodium dodecyl sulfate (SDS)-polyacrylamide gels (ePAGEL HR; 2331970, ATTO) for 75 minutes at 21 mA/mV. Gels were stained with Coomassie Brilliant Blue (EzStain Aqua; 2332370, ATTO), were imaged using the iBright FL 1500 Imaging System, and were analyzed using Image J software^77^.

## DATA AVAILABILITY

The data that support the findings of this study are available from the corresponding author upon reasonable request. The input and trajectory files have been submitted to the Biological Structure Model Archive (BSM-Arc) under BSM-ID BSM00050 (https://bsma.pdbj.org/entry/50).

## SUPPLEMENTARY DATA

## ACKNOWLEDGEMENT

We thank Shoji Takada and the laboratory members of the theoretical biophysics laboratory at Kyoto University for discussions and assistance throughout this work.

## FUNDING

This work was supported by the grant from the Kyoto University Foundation (to T.T.), the grant from the Takeda Science Foundation (to T.T.), the grant from the Shimazu Science Foundation, the Grant-in-Aid for Japan Society for the Promotion of Science Fellows (22J21003; to F.N.), the grant from the Ginpuu foundation (to F.N.).

## CONFLICT OF INTEREST

The authors have no conflict of interest, financial or otherwise.

---

**Movie S1**: The representative trajectory in which the H3/H4 tetramer was deposited to the lagging strand via the Cdc45 mediated pathway.

**Movie S2**: The representative trajectory in which the H3/H4 tetramer was deposited to the leading strand via the Cdc45 mediated pathway.

**MovieS3**: The representative trajectory in which the H3/H4 tetramer was deposited to the lagging strand via the Cdc45 unmediated pathway.

**MovieS4**: The representative trajectory in which the H3/H4 tetramer was deposited to the leading strand via the Cdc45 unmediated pathway.

**Supplementary Table S1:**
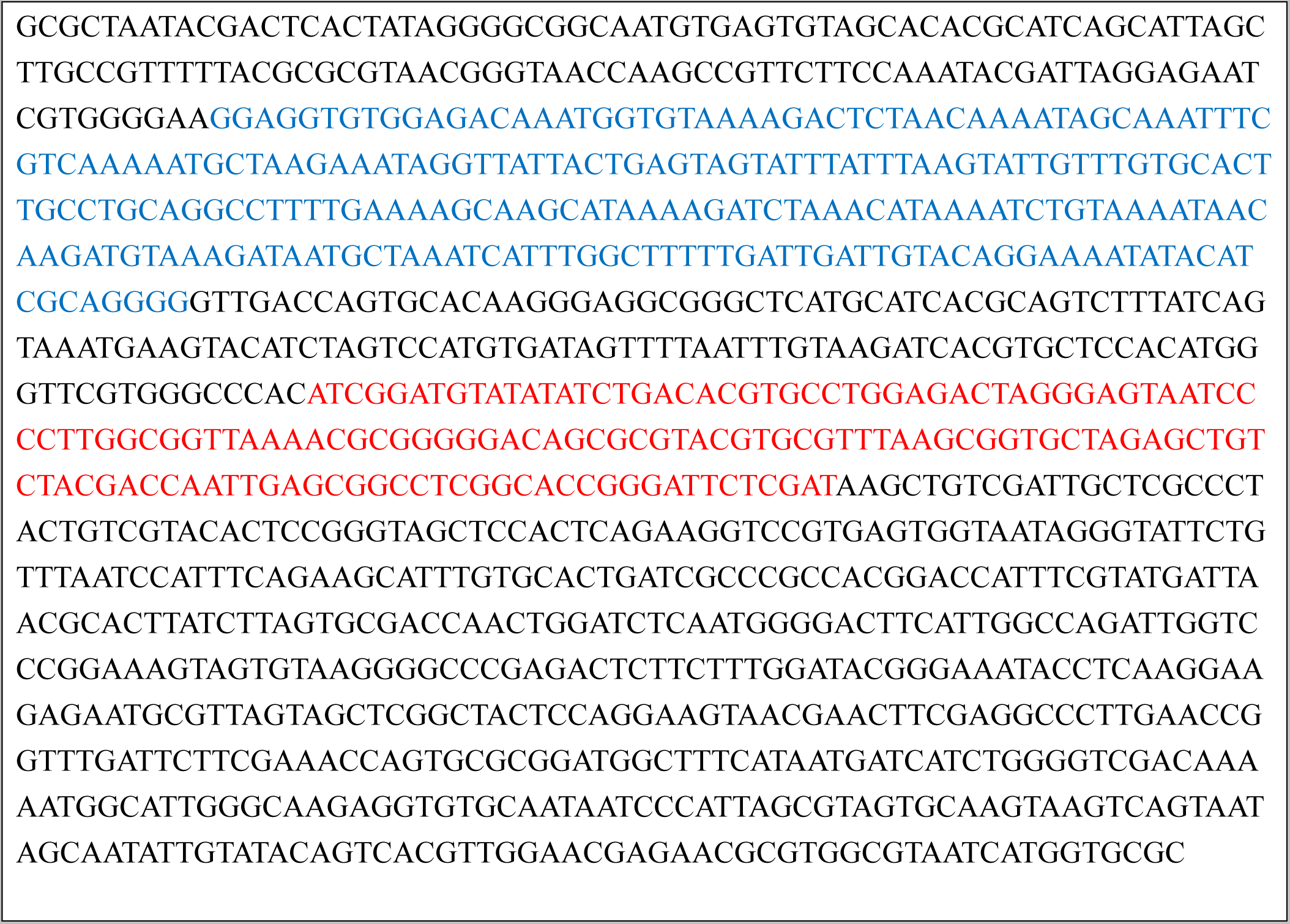
The DNA sequence used in experiments. The Autonomously Replicating Sequence 1 (ARS1) and the Widom 601 sequence are colored blue and red, respectively.

**Supplementary Figure 1:**
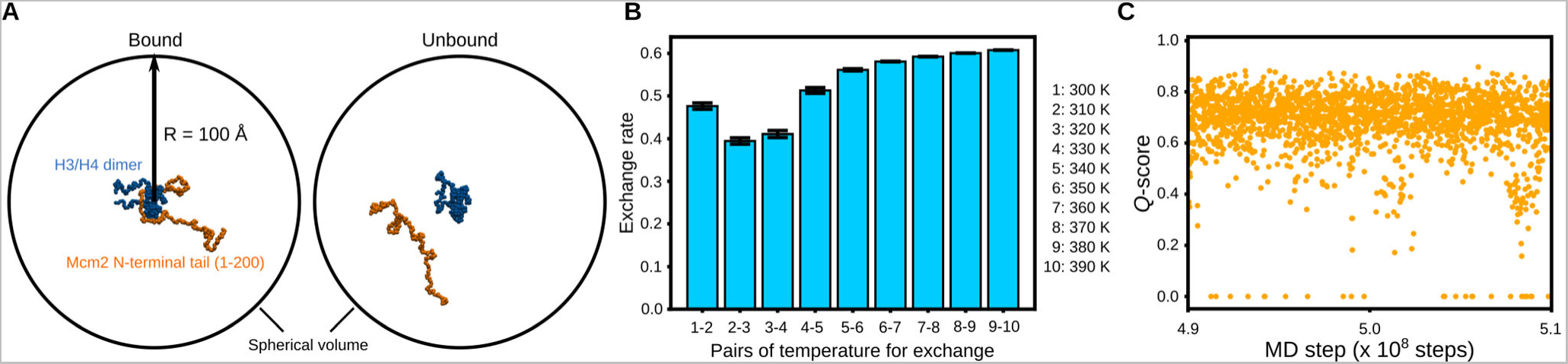
Replica exchange molecular dynamics simulations of the Mcm2 N-tail and the H3/H4 dimer in a sphere. **(A)** Representative snapshots of the bound (left) and unbound (right) states. **(B)** Exchange rates between the replicas. The standard deviations were calculated from three sets of simulations. **(C)** *Q*-scores between the Mcm2-7 N-tail and the H3/H4 dimer. The *Q*-score was defined as the ratio of residue-pair contacts at a particular time point to the natively formed contacts. We considered that a residue pair forms contact when these beads were within 1.2 times the distance in the native structure. Then, we defined the bound state as the state with *Q* > 0.

**Supplementary Figure 2:**
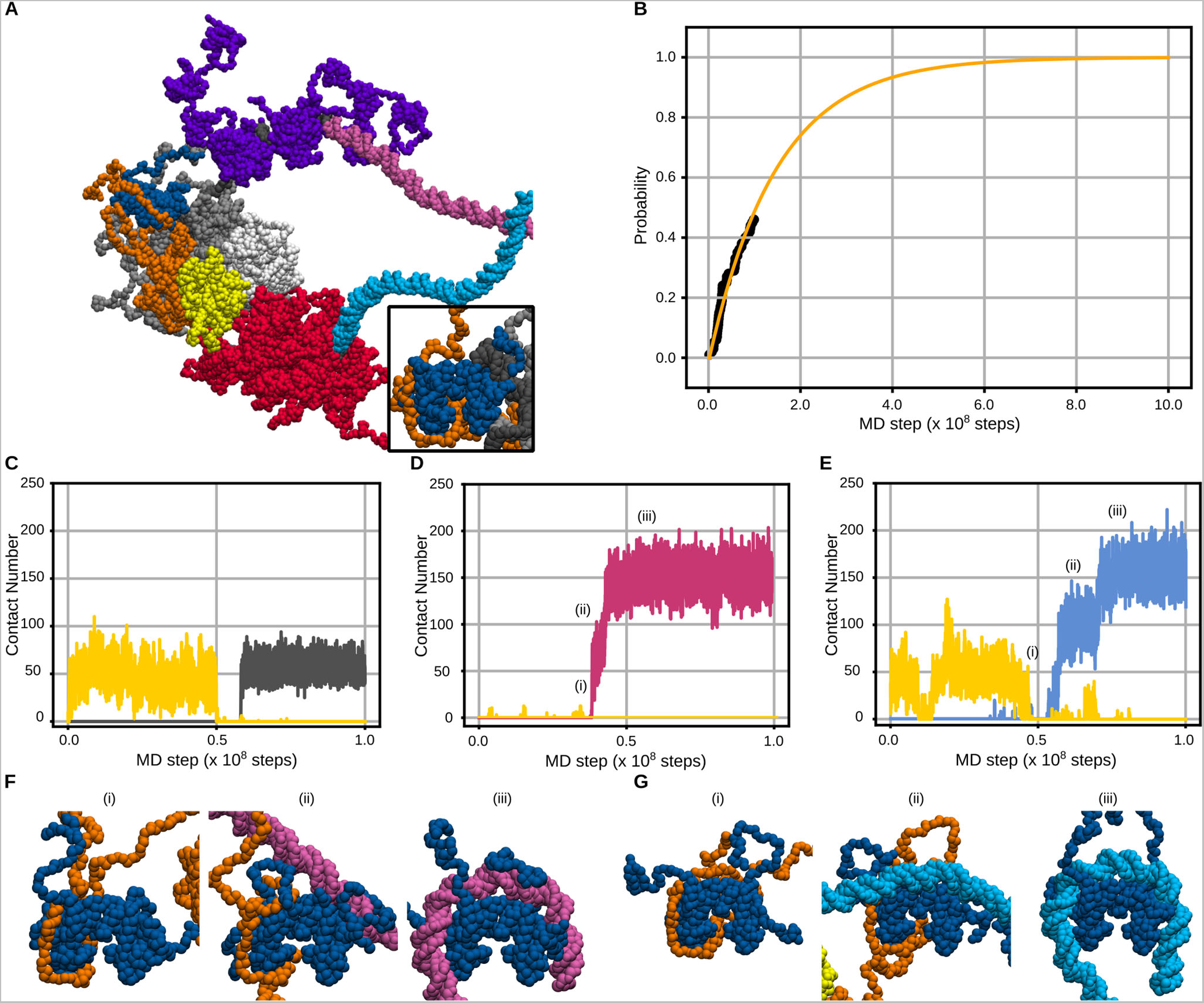
The H3/H4 tetramer was deposited on the replicated strands by the Mcm2 N-tail in the simulations of the replicated-DNA-engaged replisome. **(A)** A representative snapshot of the H3/H4 tetramer deposited on the parental strand. **(B)** Cumulative probability of the H3/H4 tetramer being deposited on the parental, lagging, and leading strands. Data points (black) and a fitting curve (yellow) are shown. **(C-E)** Time trajectories of the number of residues in the H3/H4 tetramer contacting the parental strand (C, grey), the lagging strand (D, pink), the leading strand (E, cyan), and Cdc45 (C-E, yellow). **(F, G)** Representative snapshots of the H3/H4 tetramer recycled to the lagging (F) and leading (G) strands via the Cdc45-unmediated pathway.

**Supplementary Figure 3:**
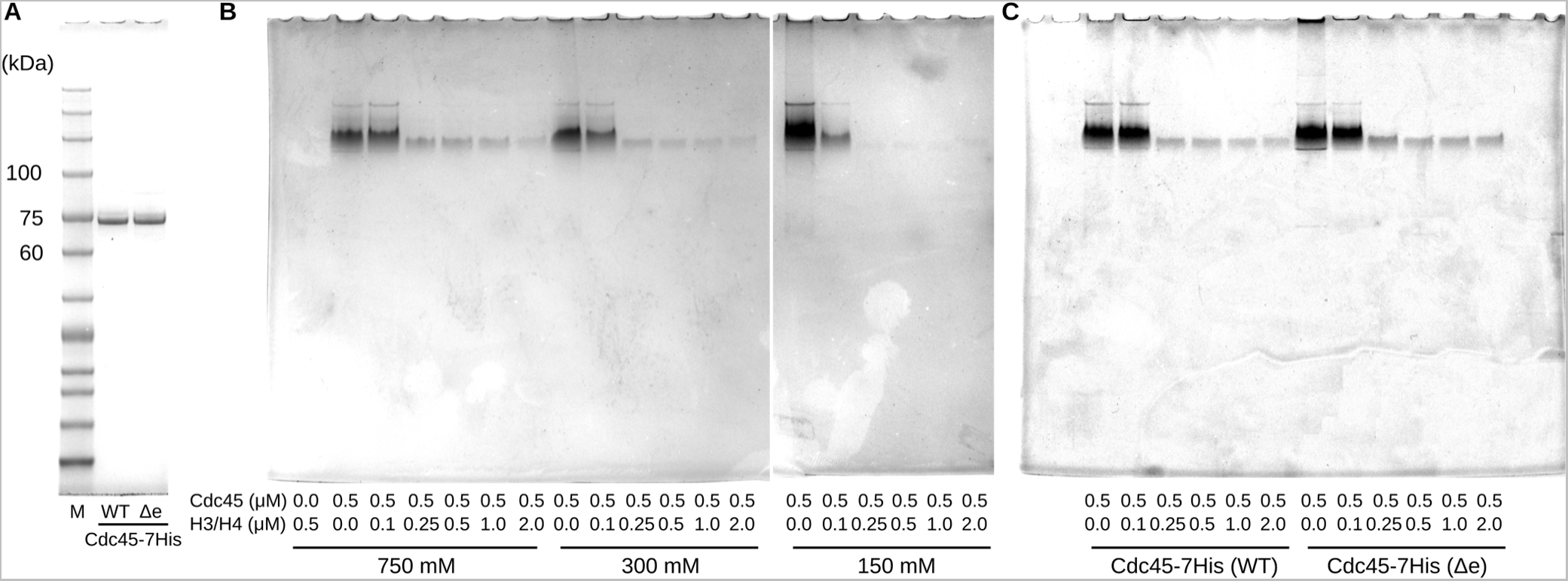
Purification of Cdc45-7His and binding assays of Cdc45 and an H3/H4 tetramer. **(A)** The gel image of 7.5% SDS-PAGE of purified Cdc45-7His and Cdc45-7His (Δe). ‘M’ denotes a marker (Nacalai Tesque Inc.; 19593-25) **(B)** Gel images of 5–20% Native-PAGE of mixtures of Cdc45-7His and H3/H4 tetramer in 150, 300, 750 mM KCl. **(C)** A gel image of 5–20% Native-PAGE of mixtures of Cdc45-7His (or Cdc45-7His (Δe)) and H3/H4 tetramers in 300 mM KCl.

**Supplementary Figure 4:**
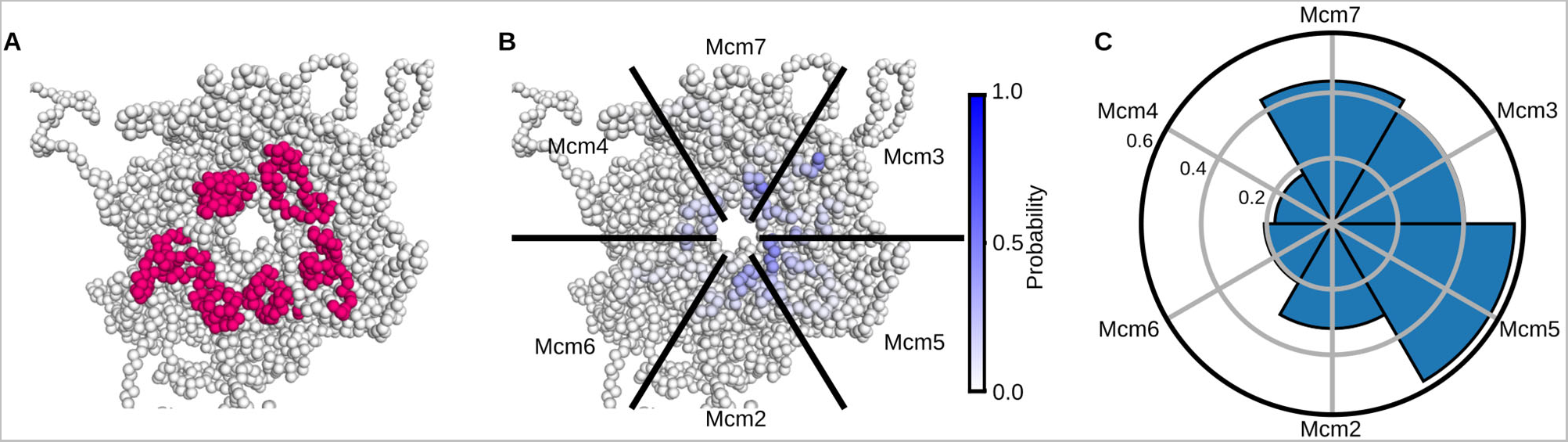
**(A & B)** Structures of Mcm2-7 viewed from the N-terminal side. The residues in zinc-finger domains of Mcm subunits were colored pink (A). The shades of blue on the structure represent the probabilities of the residues in Mcm2-7 contacting the ssDNA region of the lagging strand (B). **(C)** The probabilities of the Mcm2-7 subunits contacting the ssDNA region of the lagging strand.

**Supplementary Figure 5:**
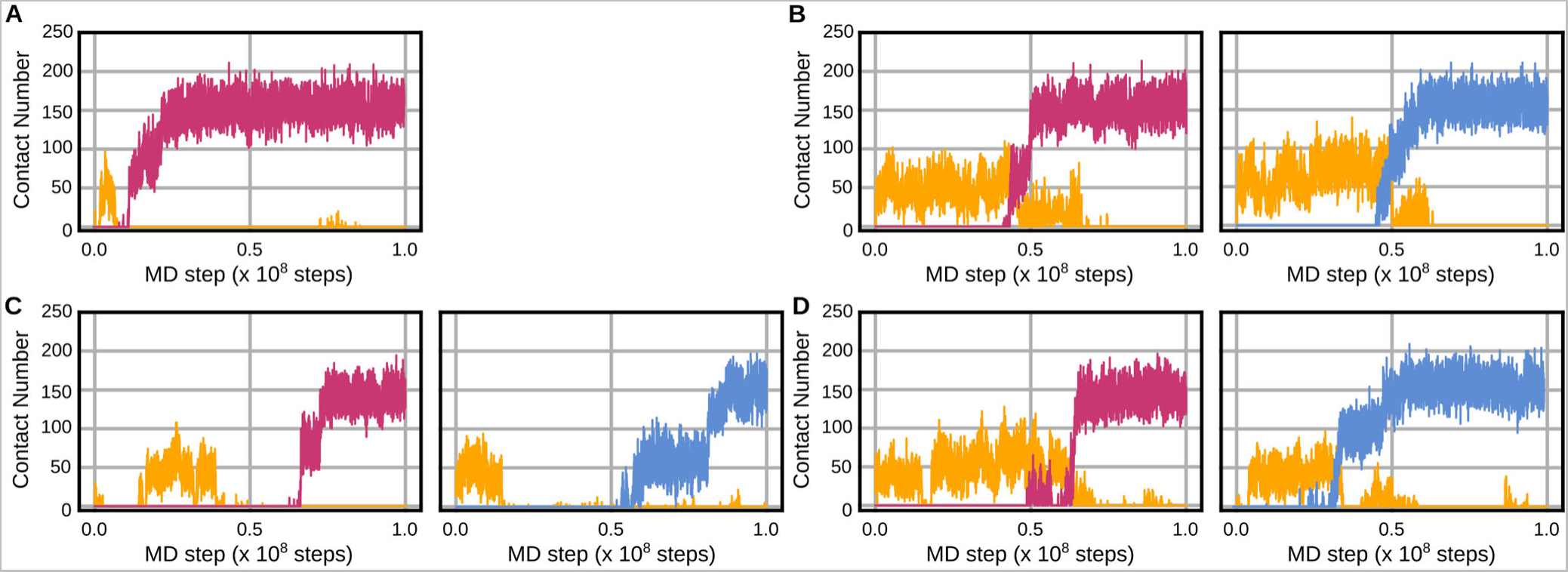
Time trajectories of the number of residues in the H3/H4 tetramer contacting the lagging strand (pink), the leading strand (cyan), and Cdc45 (yellow) in the simulations of the replicated-DNA-engaged replisome with zero or one RPA. **(A, B)** The trajectories in which the H3/H4 tetramer was recycled via the Cdc45-unmediated (A) and mediated (B) pathway in the simulations with zero RPA. **(C, D)** The trajectories in which the H3/H4 tetramer was recycled via the Cdc45-unmediated (C) and mediated (D) pathway in the simulations with one RPA.

**Supplementary Figure 6:**
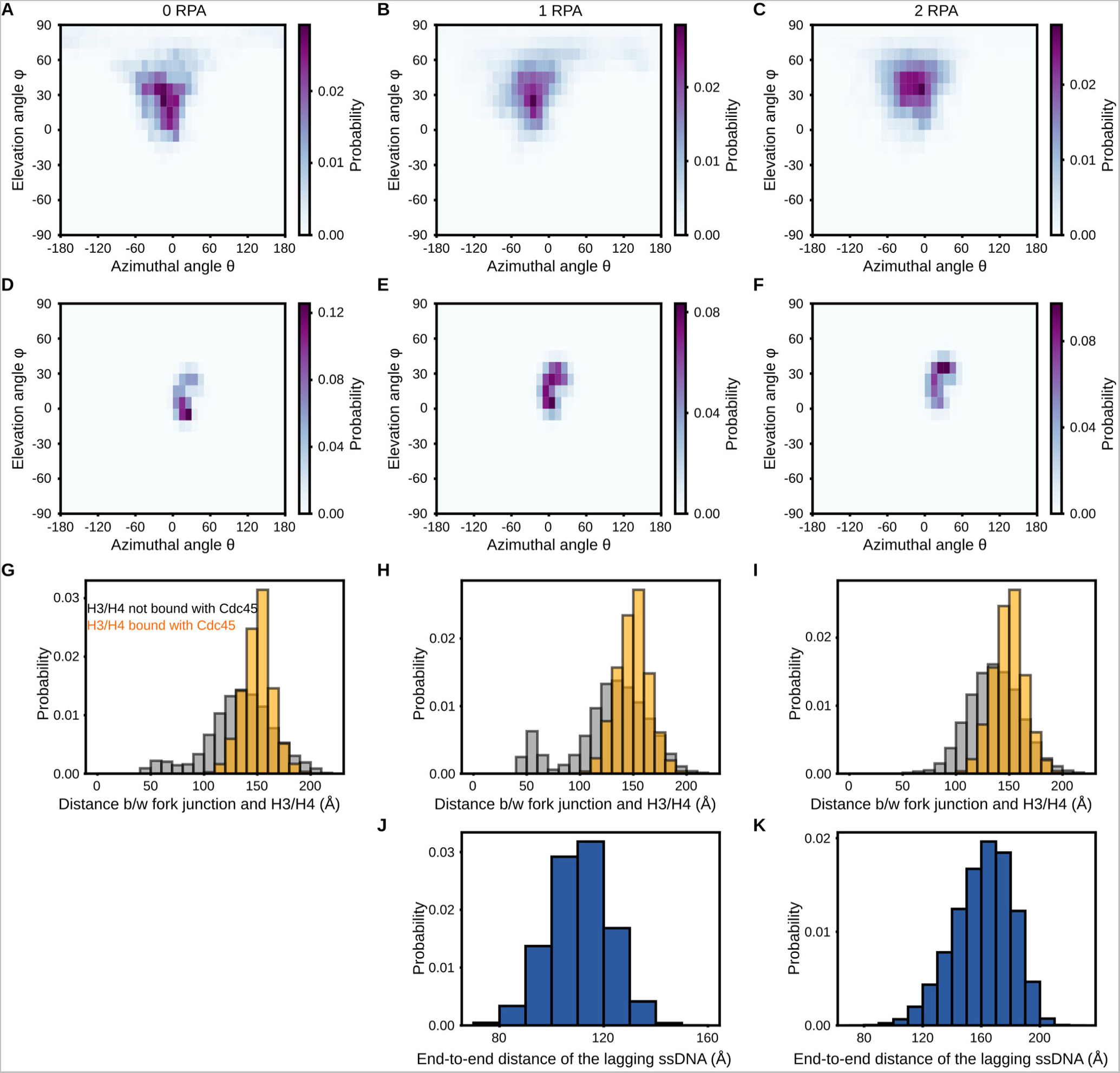
**(A-F)** 2D probability distributions of the H3/H4 tetramer not bound with Cdc45 (A, B, and C in the 0-, 1-, and 2-RPA case) and those of the H3/H4 tetramer bound with Cdc45 (D, E, and F in the 0-, 1-, and 2-RPA case) until recycling to the lagging strand. **(G-I)** 1D probability distributions of the distance between the H3/H4 tetramer bound with Cdc45 (gray) or not (orange) and the fork junction until recycling in the 0- (G), 1- (H), and 2-RPA (I) case. **(J, K)** 1D probability distributions of the end-to-end distance of the lagging ssDNA in the 1- (J) and 2-RPA (K) case.

**Supplementary Figure 7:**
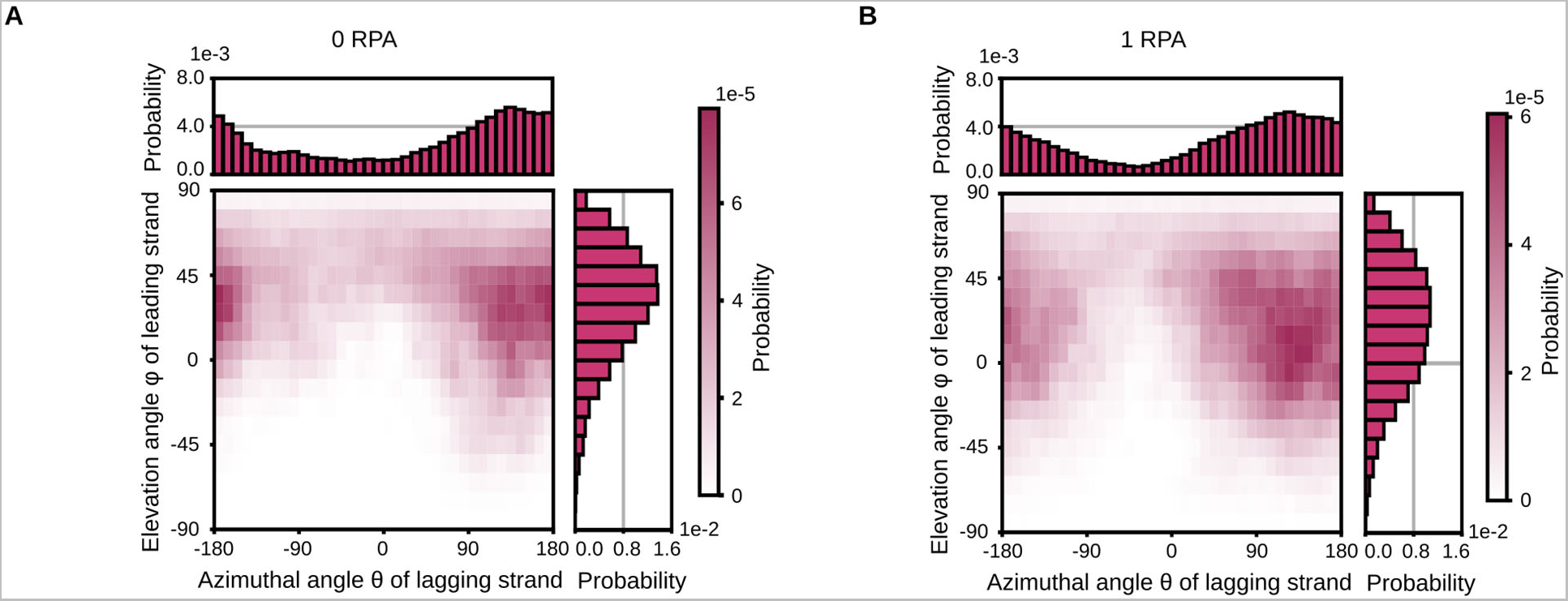
1D and 2D probability distributions of the lagging strand orientation calculated from the simulation with zero (A) and one (B) RPA molecule.

## Notes

### Competing Interest Statement

The authors have declared no competing interest.

